# Sensitivity of Bud Rot (Phytophthora sp.) Isolated from Coconut Palms in Jamaica to local Commercial Fungicide Pilarzox and Turmeric powder

**DOI:** 10.1101/2025.04.13.648621

**Authors:** Wayne Myrie, Curtis Hanchard

## Abstract

Bud rot, caused by Phytophthora sp., poses a significant threat to coconut palm production in tropical regions. This study evaluated the in vitro efficacy of the commercial fungicide Pilarzox (containing the active ingredients Cyazofamid and Cymoxanil) and turmeric (Curcuma longa) extract against Phytophthora isolates obtained from symptomatic coconut palms in St. Mary, Jamaica. Using the poisoned agar technique, three independent Phytophthora isolates were tested in five technical replicates across ten concentrations (0, 100, 150, 200, 250, 300, 500, 1,000, 5,000, and 10,000 ppm). Diagnostic tests confirmed homogeneity of variance, validating the use of ANOVA and Tukey’s HSD for statistical comparisons. Results indicated that Cyazofamid achieved complete inhibition at 500 ppm, whereas Cymoxanil required concentrations between 1,000 and 5,000 ppm for full inhibition. In contrast, turmeric exhibited a maximum inhibition of only 12% at 10,000 ppm. These findings suggest that local Phytophthora populations may have developed resistance to conventional fungicides, and while turmeric shows limited efficacy as a stand-alone agent, it may be useful as part of an integrated management strategy. Further field trials and mechanistic studies are recommended.

*(Batuman et al., 2020; Ivanov et al., 2021; Murugesh et al., 2019; Jones et al., 2023)*

## Introduction

Coconut palms (Cocos nucifera) are economically important in tropical and subtropical regions, providing resources for food, health, cosmetics, culture, and fuel (Britannica, 2024). However, bud rot—a disease predominantly caused by Phytophthora palmivora, initiates infection in the growing bud of the coconut palm. Early symptoms include water-soaked lesions that soon turn brown or black, leading to necrosis of the bud tissue (Alvarez & Peters, 2012). As the disease advances, the infection spreads from the bud to the crown, causing significant dieback and weakening of the palm, which may eventually lead to palm death if not treated early (Miller et al., 2018).

Conventional management practices include systemic fungicides; among these, Pilarzox, which contains Cyazofamid and Cymoxanil, is widely employed. Cyazofamid is a cyanoimidazole–sulphonamide fungicide (CAS 120116-88-3) used primarily to control diseases caused by oomycete pathogens. Cyazofamid inhibits mitochondrial respiration (Forkink et al., 2015) and fungal development at all stages. More specifically, it targets the Qi site of Complex III (the cytochrome bc₁ complex) in fungal mitochondria. By binding at the ubiquinone docking (Qi) site, it disrupts electron transfer within the electron transport chain and ultimately reduces ATP production. This energy deprivation stops fungal growth and development and is distinct from the modes of action of many other fungicides, helping to minimize cross-resistance (US EPA Fact Sheet for cyazofamid).

Cymoxanil is a cyanoacetamide–oxime fungicide (CAS 57966-95-7) used primarily to control downy mildew and other oomycete diseases. Although effective against oomycete pathogens, it may involve mechanisms such as dihydrofolate reductase inhibition (Kazmirchuk et al., 2024). Recent studies have shown that cymoxanil disrupts RNA synthesis by inhibiting dihydrofolate reductase (DHFR). DHFR is a key enzyme that converts dihydrofolate (DHF) into tetrahydrofolate (THF), an essential cofactor for the de novo synthesis of purines (the building blocks of RNA and DNA). Inhibition of DHFR leads to decreased THF availability, thereby reducing purine synthesis and ultimately impairing RNA production. This single-site mode of action—distinct from multi-site inhibitors—is responsible for its curative and protective effects against target pathogens (Kazmirchuk et al., 2024).

Farmers face higher expenses from fungicide applications, labour for the removal of infected material, and the costs associated with replanting or rehabilitating affected palms. For smallholder farmers, these expenses can significantly impact overall profitability (Alvarez & Peters, 2012). Recent studies have also highlighted an alarming trend of fungicide resistance in oomycete pathogens, with reports of increased resistance emerging in several regions (Jones et al., 2023; Zhang et al., 2023). This underscores the need for periodic reassessment of fungicide efficacy. Simultaneously, there is growing interest in organic alternatives, such as turmeric extract, which has demonstrated antimicrobial properties (Murugesh et al., 2019; Hernandez & Lopez, 2023). This study investigates the in vitro sensitivity of Phytophthora sp. to both Pilarzox and turmeric, aiming to inform integrated disease management strategies.

## Materials and Methods

### Pathogen Isolation and Culture

Coconut palms exhibiting late-stage bud rot were felled, and crown samples were collected from the heart region. Three independent Phytophthora isolates were obtained by culturing tissue sections on 3.9% Potato Dextrose Agar (PDA). Isolates were identified based on morphological characteristics and sub-cultured weekly to ensure purity.

### Preparation of Treatments

- **Pilarzox:** Evaluated at ten concentrations (0, 100, 150, 200, 250, 300, 500, 1,000, 5,000, and 10,000 ppm).
- **Turmeric:** Turmeric powder was dissolved (100 g in 100 mL sterile deionized water, w/v) and diluted to the same concentration range as Pilarzox.

### Poisoned Agar Technique

The method used was modified from (Ofir Degani and Gilad Cernica, 2014). After autoclaving, PDA was cooled to 55°C and amended with the appropriate concentration of either Pilarzox or turmeric. Approximately 20 mL of the amended medium was poured into 9 cm diameter petri dishes. A 10 mm agar disc from a 4–6-day-old Phytophthora colony was placed centrally in each plate. Each treatment and control (PDA without additives) were replicated five times per isolate.

### Incubation and Measurement

Plates were incubated at 28°C, and radial mycelial growth was measured along two perpendicular axes after five days. Percentage inhibition was calculated using:

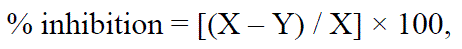

Where, X is the growth in the control and Y is the growth in treated plates.

### Statistical Analysis

Data were analysed using ANOVA followed by Tukey’s HSD post-hoc test. Homogeneity of variance was confirmed via diagnostic tests. Statistical significance was determined at p < 0.05 using IBM SPSS.

## Results

### Fungicide Efficacy

- **Cyazofamid:** A dose-dependent inhibition was observed, with complete inhibition (0 mm growth) at 500 ppm.
- **Cymoxanil:** Inhibition increased with concentration, with complete inhibition observed at 1,000– 5,000 ppm.
- **Pilarzox (Overall):** Averaging the effects of both active ingredients, effective control was achieved between 1,000 and 5,000 ppm.

### Turmeric Efficacy

Turmeric exhibited minimal inhibition, with a maximum of 12% observed at 10,000 ppm. Statistically significant differences from the control were observed at 100, 200, 5,000, and 10,000 ppm, though overall efficacy remained low.

ANOVA was used to evaluate Pilarzox’s impact on Phytophthora (see table 7). Samples were divided into nine groups based on the PPM concentration (0/control, 100, 150, 200, 250, 300, 500, 1000, 5,000, and 10,000), see table 6. Focusing on the active ingredient Cyazofamid PPM concentration. There was a statistically significant difference at the p<.05 level in satisfaction levels for the ten groups [F (9, 110) = 475.02, p=0.>0001). Focusing on the active ingredient Cymoxanil PPM concentration. There was a statistically significant difference at the p<.05 level in the ten groups [F (9, 110) = 146.820, p=0.>0001). Overall, there was a statistically significant difference at the p<.05 level in satisfaction levels for the ten groups [F (9, 110) = 117.960, p=0.>0001).

**Table 1.** Fungicide inhibition of phytophthora using Pilarzox at various Cyazofamid concentrations. *Active ingredient Cyazofamid* Post-hoc tests using the Tukey HSD (see Table 8) indicated that the PPM concentration of 0/control (M=4.207, SD=0.3433) was significantly different from all concentrations. 100 PPM Cyazofamid (M=2.540, SD=0.2221) is also significantly different from all concentrations (see table 6). At 150 PPM Cyazofamid (M=1.530, SD=0.1829) there is a significant difference compared to all samples except for at 200 PPM which shows there is no significant difference (sig=1.000) in inhibition between these two concentrations. At 200 PPM Cyazofamid (M=1.460, SD=0.2633) there is a significant difference compared to all samples except for at 150 PPM which shows there is no significant difference (sig=1.000) in inhibition between these two concentrations. At 250 PPM Cyazofamid (M=0.830, SD=0.5376) there is a significant difference compared to all samples except for at 300 PPM which shows there is no significant difference (sig=0.381) in inhibition between these two concentrations. At 300 PPM Cyazofamid (M=0.540, SD=0.3596) there is a significant difference compared to all samples except for at 250 PPM which shows there is no significant difference (sig=0.381) in inhibition between these two concentrations. At 500 PPM Cyazofamid (M=0.000, SD=0.0000) there is a significant difference compared to all samples except for at 1,000, 5,000, and 10,000 PPM which shows there is no significant difference (sig=1.000) in inhibition between these concentrations. At 1,000 PPM Cyazofamid (M=0.000, SD=0.0000), there is a significant difference compared to all samples except for at 500 and 5,000 PPM which shows there is no significant difference (sig=1.000) in inhibition between these concentrations. At 5,000 PPM Cyazofamid (M=0.000, SD=0.0000) there is a significant difference when compared to all samples except for at 500, 1,000, and 10,000 PPM which shows there is no significant difference (sig=1.000) in inhibition between these concentrations. At 10,000 PPM Cyazofamid (M=0.000, SD=0.0000) there is a significant difference compared to all samples except for at 500, 1,000, and 5,000 PPM which shows there is no significant difference (sig=1.000) in inhibition between these concentrations.

| PPM | ACTIVE INGREDIENT | % INHIBITION |
| --- | --- | --- |
| 0 | None | Control |
| 100 | *Cyazofamid* | 38.9% |
| 150 | *Cyazofamid* | 63.2% |
| 200 | *Cyazofamid* | 64.9% |
| 250 | *Cyazofamid* | 80.0% |
| 300 | *Cyazofamid* | 87.0% |
| 500 | *Cyazofamid* | 100.0% |
| 1000 | *Cyazofamid* | 100.0% |
| 5,000 | *Cyazofamid* | 100.0% |
| 10,000 | *Cyazofamid* | 100.0% |

**Table 2.**
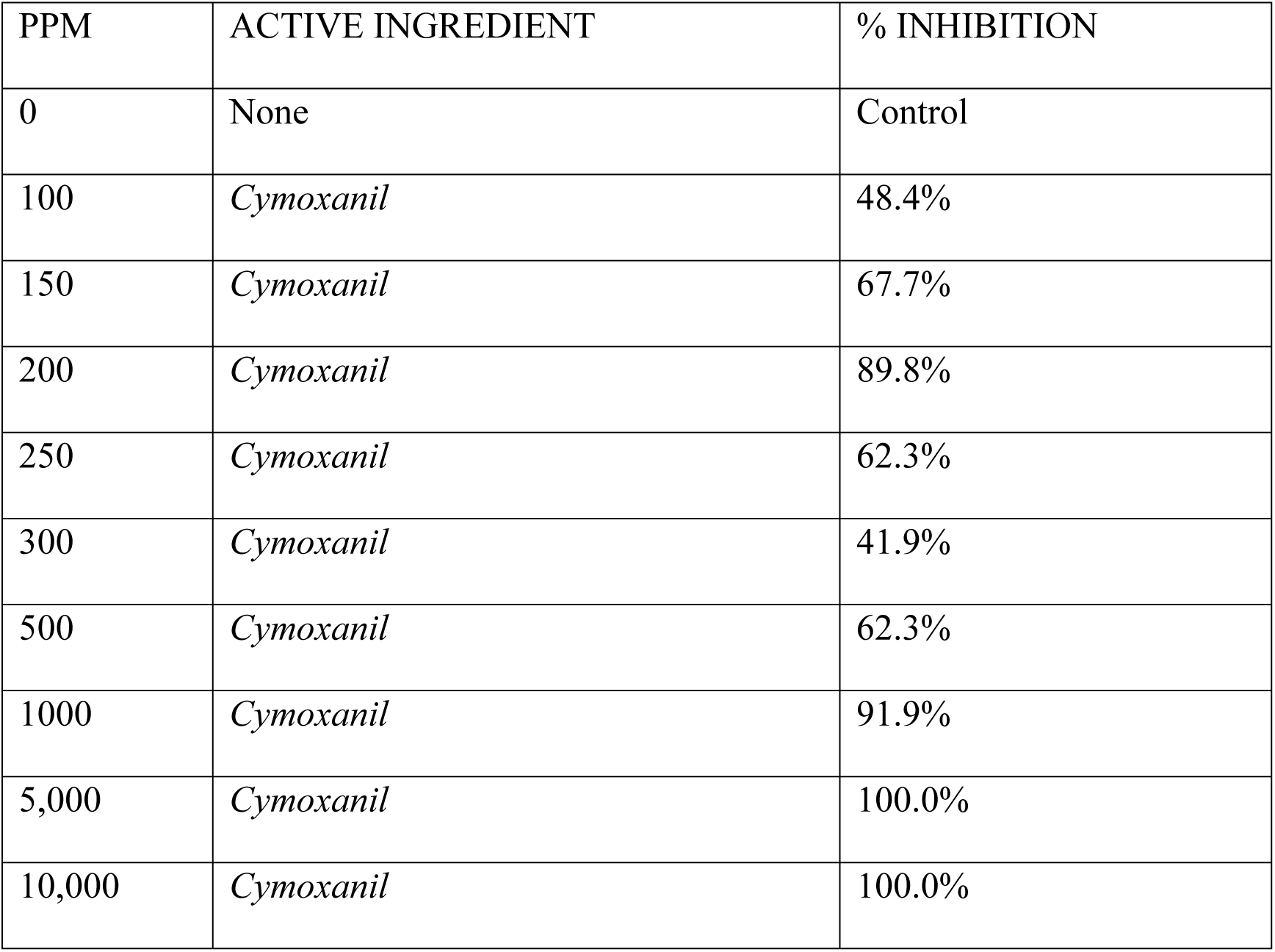
Fungicide inhibition of phytophthora using Pilarzox at various Cymoxanil concentrations. *Active ingredient Cymoxanil* Post-hoc tests using the Tukey HSD (see Table5,6,7,8,9,10,11, 12) indicated that the PPM concentration of 0/control (M=4.207, SD=0.3433) was significantly different from all concentrations. At 100 PPM Cymoxanil (M=2.220, SD=0. 72388) there is a significant difference compared to all samples, except for 250, 300, and 500 PPM which shows there is no significant difference (sig=0.123,0.940,0.123) in inhibition between these two concentrations (see table 11). At 150 PPM Cymoxanil (M=1.3900, SD=1.07750) there is a significant difference compared to all samples except for at 250 and 500 PPM which shows there is no significant difference (sig=0.983,0.983) in inhibition between these two concentrations. At 200 PPM Cymoxanil (M=.04400, SD=0.06992) there is a significant compared to all samples except for at 1000, 5,000, and 10,000 PPM which shows there is no significant difference (sig=1.000,0.521,0.521) in inhibition of these concentrations. At 250 PPM Cymoxanil (M=1.6200, SD=.69730) there is a significant difference compared to all samples except for at 100, 150 and 500 PPM which shows there is no significant difference (sig=0.123,0.983,1.000) in inhibition of these concentrations. At 300 PPM Cymoxanil (M=2.5,000, SD=0. 22111) there is a significant difference compared to all samples except for at 100 PPM which shows there is no significant difference (sig=0.940) in inhibition between these two concentrations. At 500 PPM Cymoxanil (M=1.6200, SD=0. 19322) there is a significant difference compared to all samples except for at 100, 150, and 250 PPM which shows there is no significant difference (sig=0.123,0.9831.000) in inhibition of these two concentrations. At 1,000 PPM Cymoxanil (M=0. 3500, SD=0. 05270) there is a significant difference when compared to all samples except for at 200, 5,000, and 10,000 PPM which shows there is no significant difference (sig=1.000,0.803,0.803) in inhibition. At 5,000 PPM Cymoxanil (M=0.000, SD=0.0000) there is a significant difference among all samples except for at 200, 1000, and 10,000 PPM which shows there is no significant difference (sig=0.521,0.803,1.000) in inhibition. At 10,000 PPM Cymoxanil (M=0.000, SD=0.0000) there is a significant difference among all samples except for at 200, 1,000, and 5,000 PPM which shows there is no significant difference (sig=0.521,0.803,1.000) in inhibition.

| PPM | ACTIVE INGREDIENT | % INHIBITION |
| --- | --- | --- |
| 0 | None | Control |
| 100 | *Cymoxanil* | 48.4% |
| 150 | *Cymoxanil* | 67.7% |
| 200 | *Cymoxanil* | 89.8% |
| 250 | *Cymoxanil* | 62.3% |
| 300 | *Cymoxanil* | 41.9% |
| 500 | *Cymoxanil* | 62.3% |
| 1000 | *Cymoxanil* | 91.9% |
| 5,000 | *Cymoxanil* | 100.0% |
| 10,000 | *Cymoxanil* | 100.0% |

**Table 3.** Overall fungicide inhibition of Phytophthora using Pilarzox. *Average Inhibition of Pilarzox fungicide* Post-hoc tests using the Tukey HSD (see table 14, 15, 17) indicated that the PPM concentration of 0/control (M=4.207, SD=0.3433) was significantly different from all concentrations (see table 16). 100 PPM Pilarzox (M=2.380, SD=0. 54638) there is a significant difference among all samples (sig=>0.0001). 150 PPM Pilarzox (M=1.4600, SD=0.75561) there is a significant difference among all samples except for at 200, 250, and 300 PPM which shows there is no significant difference (sig=0.177,0.963,1.000) in inhibition between these two concentrations. 200 PPM Pilarzox (M=0.9500, SD=0. 55583) there is a significant difference among all samples except for at 150,250, 300, and 500 PPM which shows there is no significant difference (sig=0.177,0.905,0.080.0.999) in inhibition between these two concentrations. 250 PPM Pilarzox (M=1.2250, SD=.72900) there is a significant difference among all samples except for 150, 200, 300, and 500 PPM which shows there is no significant difference (sig=0.963,0.905,0.862,0.457) in inhibition between these two concentrations. 300 PPM Pilarzox (M=1.5200, SD=1.04660) there is a significant difference among all samples except for at 150, 200, and 250 PPM, which shows there is no significant difference (sig=1.000,0.080,0.862) in inhibition between these two concentrations. 500 PPM Pilarzox (M=0.8100, SD=0.84161) there is a significant difference among all samples except for at 200 and 250 PPM which shows there is no significant difference (sig=0.999,0.457) in inhibition between these two concentrations. 1,000 PPM Pilarzox (M=0.1750, SD=0.18317) there is a significant difference among all samples except for at 5,000 and 10,000 PPM which shows there is no significant difference (sig=0.995,0.995) in inhibition between these two concentrations. 5,000 PPM Pilarzox (M=0.000, SD=0.0000) there is a significant difference among all samples except for at 1000 and 10,000 PPM which shows there is no significant difference (sig=0.995,1.000) in inhibition between these two concentrations. 10,000 PPM Pilarzox (M=0.000, SD=0.0000) there is a significant difference among all samples except for at 1000 and 5,000 PPM which shows no significant difference (sig=0.995,1.000) in inhibition between these two concentrations. Complete inhibition was observed at 5000 ppm (see table 3)

| PPM | AVERAGE % INHIBITION BY PILARZOX |
| --- | --- |
| 0 | 0% |
| 100 | 43.7% |
| 150 | 65.4% |
| 200 | 77.3% |
| 250 | 71.2% |
| 300 | 64.4% |
| 500 | 81.2% |
| 1000 | 95.9% |
| 5,000 | 100.0% |
| 10,000 | 100.0% |

## TURMERIC

**Table 4.** Fungicide Inhibition of Phytophthora Using Turmeric at Various Concentrations. ANOVA was conducted to evaluate the impact Turmeric (see table 19,20) has on the inhibition of phytophthora. Samples were divided into nine groups based on the PPM concentration used (0/control, 100, 150, 200, 250, 300, 500, 1000, 5,000, and 10,000). There was a statistically significant difference at the p<.05 level in satisfaction levels for the ten groups [F (9, 120) = 5.895, p=0.>001). Post-hoc tests using the Tukey HSD (see table 22) indicated that the PPM concentration of 0/control (M=4.110, SD=0.37059) was significantly different to only 100, 200, 5,000, and 10,000 (sig=0.026, 0.001, 0.008, 0.001) see table 21. 100 PPM Turmeric (M=3.6500, SD=0. 33400) there was only a significant difference at 0 PPM (sig=0.026). 150 PPM Turmeric (M=3.8400, SD=0. 33400) there is no significant difference seen amongst samples at this concentration. 200 PPM Turmeric (M=3.4900, SD=0.80753) there was only a significant difference at 0 PPM (sig=0.001). 250 PPM Turmeric (M=3.7000, SD=.32998) there is no significant difference seen amongst samples at this concentration. 300 PPM Turmeric (M=3.8100, SD=0.15239) there is no significant difference seen amongst samples at this concentration. 500 PPM Turmeric (M=3.7900, SD=0.27669) there is no significant difference seen amongst samples at this concentration. 1000 PPM Turmeric (M=3.7800, SD=0.19322), there is no significant difference seen amongst samples at this concentration. 5,000 PPM Turmeric (M=3.6000, SD=0.20548) there was only a significant difference at 0 PPM (sig=0.008). 10,000 PPM Turmeric (M=3.3500, SD=0.27183) there was only a significant difference at 0 PPM (sig=0.001). Figures (provided as supplementary material). *(Degani & Cernica, 2014; Sharadraj & Mohanan, 2014; Khan et al., 2022)*

| PPM CONCENTRATION | % INHIBITION |
| --- | --- |
| 0 | Control |
| 100 | 4% |
| 150 | -1% |
| 200 | 9% |
| 250 | 3% |
| 300 | 0% |
| 500 | 1% |
| 1000 | 1% |
| 5,000 | 6% |
| 10,000 | 12% |

## Discussion

Chart 1 shows there was a significant decrease in phytophthora growth as the concentration of Cyazofamid increases. After 500 PPM there is a plateau; thus, indicating that this is the lowest optimal PPM for the active ingredient Cyazofamid in Pilarzox in the inhibition of Phytophthora. However, it took a higher concentration of 1000 – 5,000 PPM to achieve the same effect, focusing on the active ingredient Cymoxanil (see table 1). The average effective PPM of Pilarzox was observed to be between 1000 – 5,000 PM for Pilarzox.

According to Ishihara Sangyo Kaisha Ltd, which developed Cyazofamid, its mode of action is respiratory inhibition especially at Complex Ⅲ (CIII) in the mitochondria of Oomycetes. Cyazofamid inhibits Qi (Quinone inside reducing site) of Complex Ⅲ of the said oomycetes. Complex III alongside complex II is considered the most important contributor to mitochondrial reactive oxygen species production in intact cells (Forkink et al, 2015). Cyazofamid is generally applied at 80-100 g active ingredient/Ha or 50-100 ppm concentration. In vitro testing using Pilarzox showed a 38.9% inhibition at 100 ppm active ingredient on Phytophthora sp. There was a gradual increase in inhibition by Cyazofamid with complete inhibition observed at 500 ppm after which there is a plateau. Therefore, complete inhibition is achieved at approximately 500 ppm of the active ingredient Cyazofamid. The difference between the result observed (∼500 ppm for complete inhibition) and the result using the recommended concentration of Cyazofamid (50-100 ppm), may be due to Phytophthora developing resistance to Cyazofamid. Ivanov et al, 2021 observed that continuous mass application of fungicides causes increased evolutionary pressure on *P. infestans* and consequently initiates rapid adaptation and acquisition of resistance to the fungicide involved. Cyazofamid is present in a few commercial fungicides, therefore the resistance may be present in the local population. Thus, a higher concentration of Cyazofamid is required to achieve the desired effect. Focusing on Cymoxanil, it was observed that a very high concentration of 1000 – 5,000 ppm was required to achieve complete inhibition of Phytophthora. Pilarzox optimum average inhibition was observed to be 1000-5,000 ppm. Observed results suggest the possible development of resistance of the local Phytophthora sp. to Cyazofamid and Cymoxanil-based fungicides in the field.

Cymoxanil is commonly used in the treatment of phytophthora such as *P. Infestans*. Despite its widespread use in agriculture, the exact mechanism of action remains unclear (Kazmirchuk et al, 2024). A study by Kazmirchuk et al, 2024 suggests that the Cymoxanil mode of action is the inhibition of dihydrofolate reductase enzyme activity in a dose-dependent relationship. It was observed that for complete inhibition focusing on the active ingredient Cymoxanil, high inhibition was observed at 200 ppm of 77%. This increases up to 81% at 500 ppm. Complete inhibition is only observed at 1000-5,000 ppm (see Table 2). Therefore, Phytophthora may also be experiencing increased resistance to Cymoxanil as well.

Some *Phytophthora* isolates exhibit increased expression of efflux transporters or enhanced metabolic detoxification. These mechanisms help remove or neutralize the fungicide, allowing the pathogen to survive even in the presence of chemical treatments (Pasteris et al., 2000). The resistance observed may also be associated with mutations in genes encoding the target enzymes of fungicides. For instance, alterations in the amino acid sequence of key enzymes can reduce the binding affinity of phenylamide fungicides, thereby diminishing their efficacy. As well as changes in the metabolic pathways of *Phytophthora* may also contribute to fungicide resistance. Such alterations can enable the pathogen to bypass the inhibitory effects of fungicides, ensuring continued growth and reproduction (Gisi & Sierotzki, 2008).

*Curcuma longa,* commonly called Turmeric is a tuberous rhizome known for its antioxidant and antimicrobial properties of its naturally occurring phenolic compounds (Murugesh et al, 2019). It’s believed that turmeric extract inhibits fungal growth by alteration in the morphology of the hyphae which may appear severely collapsed, plasma membrane disruption, mitochondrial destruction, lack of cytoplasm, folding of the nuclear membrane and thickened cell wall caused by chemical components of the spice extract (Murugesh et al, 2019). Martins et al 2009 study showed that Turmeric could inhibit the growth of a wide variety of fungal species. While Chen et al. (2018) identified several active compounds that contribute to its antifungal activity through multiple mechanisms. The research identified Curdione as one of turmeric’s most potent antifungal agents. It has demonstrated strong activity against pathogens such as *Fusarium graminearum*, a fungus responsible for significant crop diseases. Curcumin, the most well-known compound in turmeric, demonstrated higher efficacy than conventional antifungal agents like fluconazole against pathogens such as *Paracoccidioides brasiliensis*, which causes paracoccidioidomycosis. Curcumin can damage the fungal cell membrane, impairing its function and leading to cell death. β-elemene contributes to the overall antifungal effectiveness of turmeric extracts, reinforcing the effects of curdione and curcumin. Turmeric essential oil contains several volatile compounds, including eucalyptol, β-pinene, and camphor, which have been found to inhibit the growth of fungi such as *Aspergillus flavus*, a common contaminant in food products. Turmeric’s antifungal activity operates through multiple mechanisms, one of which is the disruption of mitochondrial function. Compounds in turmeric can interfere with ATP generation in fungal cells by targeting key mitochondrial enzymes such as succinate dehydrogenase (SDH) and NADH oxidase. This energy depletion inhibits fungal growth (Oza et al., 2021).

After the assessment of Turmeric on isolated phytophthora sp., it was proven that there was a significant difference between samples and control at 100, 200, 5,000 and 10,000 ppm (see table 19,20,21,22). The greatest inhibition observed was 12% at 10,000 ppm of Turmeric extract (see table 4). This may be because dry turmeric powder was used which may have affected the curcuminoids in the extract.

Turmeric as a stand-alone fungicide shows low effectiveness against Phytophthora. However, as an organic fungicidal cocktail, it may improve its effectiveness. Alternatively, fresh Turmeric may have significant findings compared to the Turmeric powder used. Further research is needed for Turmeric as an organic fungicide as it shows promise as a growth inhibitor of Phytophthora.

## Conclusion

This study demonstrates that while Cyazofamid is highly effective against Phytophthora sp. at 500 ppm, the significantly higher doses required for Cymoxanil suggest potential resistance issues. Although turmeric extract shows limited efficacy as a sole agent, its role in an integrated management strategy merits further exploration. These results emphasize the need for updated fungicide protocols and combined treatment approaches in managing coconut bud rot.

## Appendices Detailed tables and statistical analyses (ANOVA summaries, Tukey’s HSD comparisons) are provided as supplementary material.

## S1 Appendix Pilarzox results

**Table 5:** Analysis of Cyazofamid using SPSS.

| Cyazofamid |  |  |  |  |  |  |  |  |
| --- | --- | --- | --- | --- | --- | --- | --- | --- |
| Radial Growth |  |  |  |  |  |  |  |  |
| Concentration<br>PPM | N | Mean | Std.<br>Deviation | Std.<br>Error | 95% Confidence<br>Interval for Mean |  | Minimum | Maximum |
|  |  |  |  |  | Lower<br>Bound | Upper<br>Bound |  |  |
| 0 | 30 | 4.207 | .3433 | .0627 | 4.078 | 4.335 | 3.3 | 4.7 |
| 100 | 10 | 2.540 | .2221 | .0702 | 2.381 | 2.699 | 2.1 | 2.8 |
| 150 | 10 | 1.530 | .1829 | .0578 | 1.399 | 1.661 | 1.1 | 1.7 |
| 200 | 10 | 1.460 | .2633 | .0833 | 1.272 | 1.648 | 1.2 | 2.0 |
| 250 | 10 | .830 | .5376 | .1700 | .445 | 1.215 | .0 | 1.5 |
| 300 | 10 | .540 | .3596 | .1137 | .283 | .797 | .0 | 1.2 |
| 500 | 10 | .000 | .0000 | .0000 | .000 | .000 | .0 | .0 |
| 1000 | 10 | .000 | .0000 | .0000 | .000 | .000 | .0 | .0 |
| 5,000 | 10 | .000 | .0000 | .0000 | .000 | .000 | .0 | .0 |
| 10,000 | 10 | .000 | .0000 | .0000 | .000 | .000 | .0 | .0 |
| Total | 120 | 1.627 | 1.6927 | .1545 | 1.321 | 1.933 | .0 | 4.7 |

**Table 6:** Multiple Comparison Analysis of Cyazofamid using SPSS.

| Multiple Comparisons for Cyazofamid |  |  |  |  |  |  |
| --- | --- | --- | --- | --- | --- | --- |
| Dependent Variable: Radial Growth |  |  |  |  |  |  |
| Tukey HSD |  |  |  |  |  |  |
| (I) PPM Concentration | (J) PPM Concentration | Mean Difference (I-J) | Std. Error | Sig. | 95% Confidence Interval |  |
|  |  |  |  |  | Lower Bound | Upper Bound |
| 0 | 100 | 1.6667* | .1018 | .000 | 1.338 | 1.995 |
|  | 150 | 2.6767* | .1018 | .000 | 2.348 | 3.005 |
|  | 200 | 2.7467* | .1018 | .000 | 2.418 | 3.075 |
|  | 250 | 3.3767* | .1018 | .000 | 3.048 | 3.705 |
|  | 300 | 3.6667* | .1018 | .000 | 3.338 | 3.995 |
|  | 500 | 4.2067* | .1018 | .000 | 3.878 | 4.535 |
|  | 1000 | 4.2067* | .1018 | .000 | 3.878 | 4.535 |
|  | 5,000 | 4.2067* | .1018 | .000 | 3.878 | 4.535 |
|  | 10,000 | 4.2067* | .1018 | .000 | 3.878 | 4.535 |
| 100 | 0 | -1.6667* | .1018 | .000 | -1.995 | -1.338 |
|  | 150 | 1.0100* | .1247 | .000 | .607 | 1.413 |
|  | 200 | 1.0800* | .1247 | .000 | .677 | 1.483 |
|  | 250 | 1.7100* | .1247 | .000 | 1.307 | 2.113 |
|  | 300 | 2.0000* | .1247 | .000 | 1.597 | 2.403 |
|  | 500 | 2.5400* | .1247 | .000 | 2.137 | 2.943 |
|  | 1000 | 2.5400* | .1247 | .000 | 2.137 | 2.943 |
|  | 5,000 | 2.5400* | .1247 | .000 | 2.137 | 2.943 |
|  | 10,000 | 2.5400* | .1247 | .000 | 2.137 | 2.943 |
| 150 | 0 | -2.6767* | .1018 | .000 | -3.005 | -2.348 |
|  | 100 | -1.0100* | .1247 | .000 | -1.413 | -.607 |
|  | 200 | .0700 | .1247 | 1.000 | -.333 | .473 |
|  | 250 | .7000* | .1247 | .000 | .297 | 1.103 |
|  | 300 | .9900* | .1247 | .000 | .587 | 1.393 |
|  | 500 | 1.5300* | .1247 | .000 | 1.127 | 1.933 |
|  | 1000 | 1.5300* | .1247 | .000 | 1.127 | 1.933 |
|  | 5,000 | 1.5300* | .1247 | .000 | 1.127 | 1.933 |
|  | 10,000 | 1.5300* | .1247 | .000 | 1.127 | 1.933 |
| 200 | 0 | -2.7467* | .1018 | .000 | -3.075 | -2.418 |
|  | 100 | -1.0800* | .1247 | .000 | -1.483 | -.677 |
|  | 150 | -.0700 | .1247 | 1.000 | -.473 | .333 |
|  | 250 | .6300* | .1247 | .000 | .227 | 1.033 |
|  | 300 | .9200* | .1247 | .000 | .517 | 1.323 |
|  | 500 | 1.4600* | .1247 | .000 | 1.057 | 1.863 |
|  | 1000 | 1.4600* | .1247 | .000 | 1.057 | 1.863 |
|  | 5,000 | 1.4600* | .1247 | .000 | 1.057 | 1.863 |
|  | 10,000 | 1.4600* | .1247 | .000 | 1.057 | 1.863 |
| 250 | 0 | -3.3767* | .1018 | .000 | -3.705 | -3.048 |
|  | 100 | -1.7100* | .1247 | .000 | -2.113 | -1.307 |
|  | 150 | -.7000* | .1247 | .000 | -1.103 | -.297 |
|  | 200 | -.6300* | .1247 | .000 | -1.033 | -.227 |
|  | 300 | .2900 | .1247 | .381 | -.113 | .693 |
|  | 500 | .8300* | .1247 | .000 | .427 | 1.233 |
|  | 1000 | .8300* | .1247 | .000 | .427 | 1.233 |
|  | 5,000 | .8300* | .1247 | .000 | .427 | 1.233 |
|  | 10,000 | .8300* | .1247 | .000 | .427 | 1.233 |
| 300 | 0 | -3.6667* | .1018 | .000 | -3.995 | -3.338 |
|  | 100 | -2.0000* | .1247 | .000 | -2.403 | -1.597 |
|  | 150 | -.9900* | .1247 | .000 | -1.393 | -.587 |
|  | 200 | -.9200* | .1247 | .000 | -1.323 | -.517 |
|  | 250 | -.2900 | .1247 | .381 | -.693 | .113 |
|  | 500 | .5400* | .1247 | .001 | .137 | .943 |
|  | 1000 | .5400* | .1247 | .001 | .137 | .943 |
|  | 5,000 | .5400* | .1247 | .001 | .137 | .943 |
|  | 10,000 | .5400* | .1247 | .001 | .137 | .943 |
| 500 | 0 | -4.2067* | .1018 | .000 | -4.535 | -3.878 |
|  | 100 | -2.5400* | .1247 | .000 | -2.943 | -2.137 |
|  | 150 | -1.5300* | .1247 | .000 | -1.933 | -1.127 |
|  | 200 | -1.4600* | .1247 | .000 | -1.863 | -1.057 |
|  | 250 | -.8300* | .1247 | .000 | -1.233 | -.427 |
|  | 300 | -.5400* | .1247 | .001 | -.943 | -.137 |
|  | 1000 | .0000 | .1247 | 1.000 | -.403 | .403 |
|  | 5,000 | .0000 | .1247 | 1.000 | -.403 | .403 |
|  | 10,000 | .0000 | .1247 | 1.000 | -.403 | .403 |
| 1000 | 0 | -4.2067* | .1018 | .000 | -4.535 | -3.878 |
|  | 100 | -2.5400* | .1247 | .000 | -2.943 | -2.137 |
|  | 150 | -1.5300* | .1247 | .000 | -1.933 | -1.127 |
|  | 200 | -1.4600* | .1247 | .000 | -1.863 | -1.057 |
|  | 250 | -.8300* | .1247 | .000 | -1.233 | -.427 |
|  | 300 | -.5400* | .1247 | .001 | -.943 | -.137 |
|  | 500 | .0000 | .1247 | 1.000 | -.403 | .403 |
|  | 5,000 | .0000 | .1247 | 1.000 | -.403 | .403 |
|  | 10,000 | .0000 | .1247 | 1.000 | -.403 | .403 |
| 5,000 | 0 | -4.2067* | .1018 | .000 | -4.535 | -3.878 |
|  | 100 | -2.5400* | .1247 | .000 | -2.943 | -2.137 |
|  | 150 | -1.5300* | .1247 | .000 | -1.933 | -1.127 |
|  | 200 | -1.4600* | .1247 | .000 | -1.863 | -1.057 |
|  | 250 | -.8300* | .1247 | .000 | -1.233 | -.427 |
|  | 300 | -.5400* | .1247 | .001 | -.943 | -.137 |
|  | 500 | .0000 | .1247 | 1.000 | -.403 | .403 |
|  | 1000 | .0000 | .1247 | 1.000 | -.403 | .403 |
|  | 10,000 | .0000 | .1247 | 1.000 | -.403 | .403 |
| 10,000 | 0 | -4.2067* | .1018 | .000 | -4.535 | -3.878 |
|  | 100 | -2.5400* | .1247 | .000 | -2.943 | -2.137 |
|  | 150 | -1.5300* | .1247 | .000 | -1.933 | -1.127 |
|  | 200 | -1.4600* | .1247 | .000 | -1.863 | -1.057 |
|  | 250 | -.8300* | .1247 | .000 | -1.233 | -.427 |
|  | 300 | -.5400* | .1247 | .001 | -.943 | -.137 |
|  | 500 | .0000 | .1247 | 1.000 | -.403 | .403 |
|  | 1000 | .0000 | .1247 | 1.000 | -.403 | .403 |
|  | 5,000 | .0000 | .1247 | 1.000 | -.403 | .403 |

\*. The mean difference is significant at the 0.05 level.

**Table 7:** ANOVA Analysis of Cyazofamid using SPSS.

| **Cyazofamid ANOVA** |  |  |  |  |  |
| --- | --- | --- | --- | --- | --- |
| Radial Growth |  |  |  |  |  |
|  | Sum of Squares | df | Mean Square | F | Sig. |
| Between different PPM concentration | 332.402 | 9 | 36.934 | 475.020 | .000 |
| Within PPM concentrations | 8.553 | 110 | .078 |  |  |
| Total | 340.955 | 119 |  |  |  |

**Table 8:** Tukey HSD Analysis of Cyazofamid using SPSS.

| **Radial Growth of Cyazofamid** |  |  |  |  |  |  |
| --- | --- | --- | --- | --- | --- | --- |
| Tukey HSD<sup>a,b</sup> |  |  |  |  |  |  |
| PPM Concentration | N | Subset for alpha = 0.05 |  |  |  |  |
|  |  | 1 | 2 | 3 | 4 | 5 |
| 500 | 10 | .000 |  |  |  |  |
| 1000 | 10 | .000 |  |  |  |  |
| 5,000 | 10 | .000 |  |  |  |  |
| 10,000 | 10 | .000 |  |  |  |  |
| 300 | 10 |  | .540 |  |  |  |
| 250 | 10 |  | .830 |  |  |  |
| 200 | 10 |  |  | 1.460 |  |  |
| 150 | 10 |  |  | 1.530 |  |  |
| 100 | 10 |  |  |  | 2.540 |  |
| 0 | 30 |  |  |  |  | 4.207 |
| Sig. |  | 1.000 | .332 | 1.000 | 1.000 | 1.000 |
| Means for groups in homogeneous subsets are displayed. |  |  |  |  |  |  |
| a. Uses Harmonic Mean Sample Size = 10.714. |  |  |  |  |  |  |
| b. The group sizes are unequal. The harmonic mean of the group sizes is used. Type I error levels are not guaranteed. |  |  |  |  |  |  |

## S2 Appendix

**Table 9:** Analysis of Cymoxanil using SPSS.

| **Cymoxanil Descriptives** |  |  |  |  |  |  |  |  |
| --- | --- | --- | --- | --- | --- | --- | --- | --- |
| Radial Growth |  |  |  |  |  |  |  |  |
|  | N | Mean | Std.<br>Deviation | Std.<br>Error | 95% Confidence Interval<br>for Mean |  | Minimum | Maximum |
|  |  |  |  |  | Lower<br>Bound | Upper<br>Bound |  |  |
| 0 | 30 | 4.2067 | .34334 | .06269 | 4.0785 | 4.3349 | 3.30 | 4.70 |
| 100 | 10 | 2.2200 | .72388 | .22891 | 1.7022 | 2.7378 | 1.40 | 3.40 |
| 150 | 10 | 1.3900 | 1.07750 | .34073 | .6192 | 2.1608 | .30 | 2.90 |
| 200 | 10 | .4400 | .06992 | .02211 | .3900 | .4900 | .40 | .60 |
| 250 | 10 | 1.6200 | .69730 | .22050 | 1.1212 | 2.1188 | .40 | 2.40 |
| 300 | 10 | 2.5,000 | .22111 | .06992 | 2.3418 | 2.6582 | 2.10 | 2.80 |
| 500 | 10 | 1.6200 | .19322 | .06110 | 1.4818 | 1.7582 | 1.40 | 1.90 |
| 1000 | 10 | .3500 | .05270 | .01667 | .3123 | .3877 | .30 | .40 |
| 5,000 | 10 | .0000 | .00000 | .00000 | .0000 | .0000 | .00 | .00 |
| 10,000 | 10 | .0000 | .00000 | .00000 | .0000 | .0000 | .00 | .00 |
| Total | 120 | 1.8967 | 1.61339 | .14728 | 1.6050 | 2.1883 | .00 | 4.70 |

**Table 10:** ANOVA Analysis of Cymoxanil using SPSS.

| ANOVA for Cymoxanil |  |  |  |  |  |
| --- | --- | --- | --- | --- | --- |
| Radial Growth |  |  |  |  |  |
|  | Sum of Squares | df | Mean Square | F | Sig. |
| Between Groups | 285.954 | 9 | 31.773 | 146.820 | .000 |
| Within Groups | 23.805 | 110 | .216 |  |  |
| Total | 309.759 | 119 |  |  |  |

**Table 11:** Multiple Comparison Analysis of Cymoxanil using SPSS.

|  |
| --- |
| **Multiple Comparisons for Cymoxanil** |
| Dependent Variable: Radial Growth |

| Tukey HSD |  |  |  |  |  |  |
| --- | --- | --- | --- | --- | --- | --- |
| (I) PPM<br>Concentration | (J) PPM<br>Concentration | Mean<br>Difference (I-J) | Std.<br>Error | Sig. | 95% Confidence<br>Interval |  |
|  |  |  |  |  | Lower<br>Bound | Upper<br>Bound |
| 0 | 100 | 1.98667* | .16987 | .000 | 1.4381 | 2.5353 |
|  | 150 | 2.81667* | .16987 | .000 | 2.2681 | 3.3653 |
|  | 200 | 3.76667* | .16987 | .000 | 3.2181 | 4.3153 |
|  | 250 | 2.58667* | .16987 | .000 | 2.0381 | 3.1353 |
|  | 300 | 1.70667* | .16987 | .000 | 1.1581 | 2.2553 |
|  | 500 | 2.58667* | .16987 | .000 | 2.0381 | 3.1353 |
|  | 1000 | 3.85667* | .16987 | .000 | 3.3081 | 4.4053 |
|  | 5,000 | 4.20667* | .16987 | .000 | 3.6581 | 4.7553 |
|  | 10,000 | 4.20667* | .16987 | .000 | 3.6581 | 4.7553 |
| 100 | 0 | -1.98667* | .16987 | .000 | -2.5353 | -1.4381 |
|  | 150 | .83000* | .20804 | .005 | .1581 | 1.5019 |
|  | 200 | 1.78000* | .20804 | .000 | 1.1081 | 2.4519 |
|  | 250 | .60000 | .20804 | .123 | -.0719 | 1.2719 |
|  | 300 | -.28000 | .20804 | .940 | -.9519 | .3919 |
|  | 500 | .60000 | .20804 | .123 | -.0719 | 1.2719 |
|  | 1000 | 1.87000* | .20804 | .000 | 1.1981 | 2.5419 |
|  | 5,000 | 2.22000* | .20804 | .000 | 1.5481 | 2.8919 |
|  | 10,000 | 2.22000* | .20804 | .000 | 1.5481 | 2.8919 |
| 150 | 0 | -2.81667* | .16987 | .000 | -3.3653 | -2.2681 |
|  | 100 | -.83000* | .20804 | .005 | -1.5019 | -.1581 |
|  | 200 | .95,000* | .20804 | .001 | .2781 | 1.6219 |
|  | 250 | -.23000 | .20804 | .983 | -.9019 | .4419 |
|  | 300 | -1.11000* | .20804 | .000 | -1.7819 | -.4381 |
|  | 500 | -.23000 | .20804 | .983 | -.9019 | .4419 |
|  | 1000 | 1.04000* | .20804 | .000 | .3681 | 1.7119 |
|  | 5,000 | 1.39000* | .20804 | .000 | .7181 | 2.0619 |
|  | 10,000 | 1.39000* | .20804 | .000 | .7181 | 2.0619 |
| 200 | 0 | -3.76667* | .16987 | .000 | -4.3153 | -3.2181 |
|  | 100 | -1.78000* | .20804 | .000 | -2.4519 | -1.1081 |
|  | 150 | -.95,000* | .20804 | .001 | -1.6219 | -.2781 |
|  | 250 | -1.18000* | .20804 | .000 | -1.8519 | -.5081 |
|  | 300 | -2.06000* | .20804 | .000 | -2.7319 | -1.3881 |
|  | 500 | -1.18000* | .20804 | .000 | -1.8519 | -.5081 |
|  | 1000 | .09000 | .20804 | 1.000 | -.5819 | .7619 |
|  | 5,000 | .44000 | .20804 | .521 | -.2319 | 1.1119 |
|  | 10,000 | .44000 | .20804 | .521 | -.2319 | 1.1119 |
| 250 | 0 | -2.58667* | .16987 | .000 | -3.1353 | -2.0381 |
|  | 100 | -.60000 | .20804 | .123 | -1.2719 | .0719 |
|  | 150 | .23000 | .20804 | .983 | -.4419 | .9019 |
|  | 200 | 1.18000* | .20804 | .000 | .5081 | 1.8519 |
|  | 300 | -.88000* | .20804 | .002 | -1.5519 | -.2081 |
|  | 500 | .00000 | .20804 | 1.000 | -.6719 | .6719 |
|  | 1000 | 1.27000* | .20804 | .000 | .5981 | 1.9419 |
|  | 5,000 | 1.62000* | .20804 | .000 | .9481 | 2.2919 |
|  | 10,000 | 1.62000* | .20804 | .000 | .9481 | 2.2919 |
| 300 | 0 | -1.70667* | .16987 | .000 | -2.2553 | -1.1581 |
|  | 100 | .28000 | .20804 | .940 | -.3919 | .9519 |
|  | 150 | 1.11000* | .20804 | .000 | .4381 | 1.7819 |
|  | 200 | 2.06000* | .20804 | .000 | 1.3881 | 2.7319 |
|  | 250 | .88000* | .20804 | .002 | .2081 | 1.5519 |
|  | 500 | .88000* | .20804 | .002 | .2081 | 1.5519 |
|  | 1000 | 2.15,000* | .20804 | .000 | 1.4781 | 2.8219 |
|  | 5,000 | 2.5,0000* | .20804 | .000 | 1.8281 | 3.1719 |
|  | 10,000 | 2.5,0000* | .20804 | .000 | 1.8281 | 3.1719 |
| 500 | 0 | -2.58667* | .16987 | .000 | -3.1353 | -2.0381 |
|  | 100 | -.60000 | .20804 | .123 | -1.2719 | .0719 |
|  | 150 | .23000 | .20804 | .983 | -.4419 | .9019 |
|  | 200 | 1.18000* | .20804 | .000 | .5081 | 1.8519 |
|  | 250 | .00000 | .20804 | 1.000 | -.6719 | .6719 |
|  | 300 | -.88000* | .20804 | .002 | -1.5519 | -.2081 |
|  | 1000 | 1.27000* | .20804 | .000 | .5981 | 1.9419 |
|  | 5,000 | 1.62000* | .20804 | .000 | .9481 | 2.2919 |
|  | 10,000 | 1.62000* | .20804 | .000 | .9481 | 2.2919 |
| 1000 | 0 | -3.85667* | .16987 | .000 | -4.4053 | -3.3081 |
|  | 100 | -1.87000* | .20804 | .000 | -2.5419 | -1.1981 |
|  | 150 | -1.04000* | .20804 | .000 | -1.7119 | -.3681 |
|  | 200 | -.09000 | .20804 | 1.000 | -.7619 | .5819 |
|  | 250 | -1.27000* | .20804 | .000 | -1.9419 | -.5981 |
|  | 300 | -2.15,000* | .20804 | .000 | -2.8219 | -1.4781 |
|  | 500 | -1.27000* | .20804 | .000 | -1.9419 | -.5981 |
|  | 5,000 | .35,000 | .20804 | .803 | -.3219 | 1.0219 |
|  | 10,000 | .35,000 | .20804 | .803 | -.3219 | 1.0219 |
| 5,000 | 0 | -4.20667* | .16987 | .000 | -4.7553 | -3.6581 |
|  | 100 | -2.22000* | .20804 | .000 | -2.8919 | -1.5481 |
|  | 150 | -1.39000* | .20804 | .000 | -2.0619 | -.7181 |
|  | 200 | -.44000 | .20804 | .521 | -1.1119 | .2319 |
|  | 250 | -1.62000* | .20804 | .000 | -2.2919 | -.9481 |
|  | 300 | -2.5,0000* | .20804 | .000 | -3.1719 | -1.8281 |
|  | 500 | -1.62000* | .20804 | .000 | -2.2919 | -.9481 |
|  | 1000 | -.35,000 | .20804 | .803 | -1.0219 | .3219 |
|  | 10,000 | .00000 | .20804 | 1.000 | -.6719 | .6719 |
| 10,000 | 0 | -4.20667* | .16987 | .000 | -4.7553 | -3.6581 |
|  | 100 | -2.22000* | .20804 | .000 | -2.8919 | -1.5481 |
|  | 150 | -1.39000* | .20804 | .000 | -2.0619 | -.7181 |
|  | 200 | -.44000 | .20804 | .521 | -1.1119 | .2319 |
|  | 250 | -1.62000* | .20804 | .000 | -2.2919 | -.9481 |
|  | 300 | -2.5,0000* | .20804 | .000 | -3.1719 | -1.8281 |
|  | 500 | -1.62000* | .20804 | .000 | -2.2919 | -.9481 |
|  | 1000 | -.35,000 | .20804 | .803 | -1.0219 | .3219 |
|  | 5,000 | .00000 | .20804 | 1.000 | -.6719 | .6719 |

\*. The mean difference is significant at the 0.05 level.

**Table 12:**
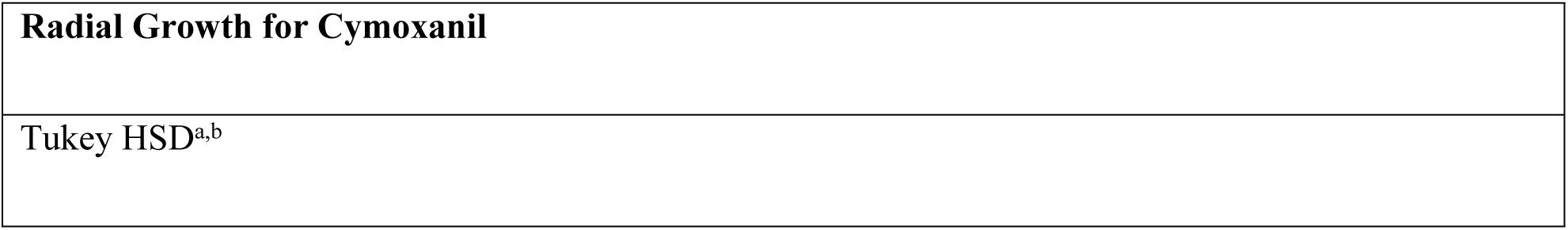
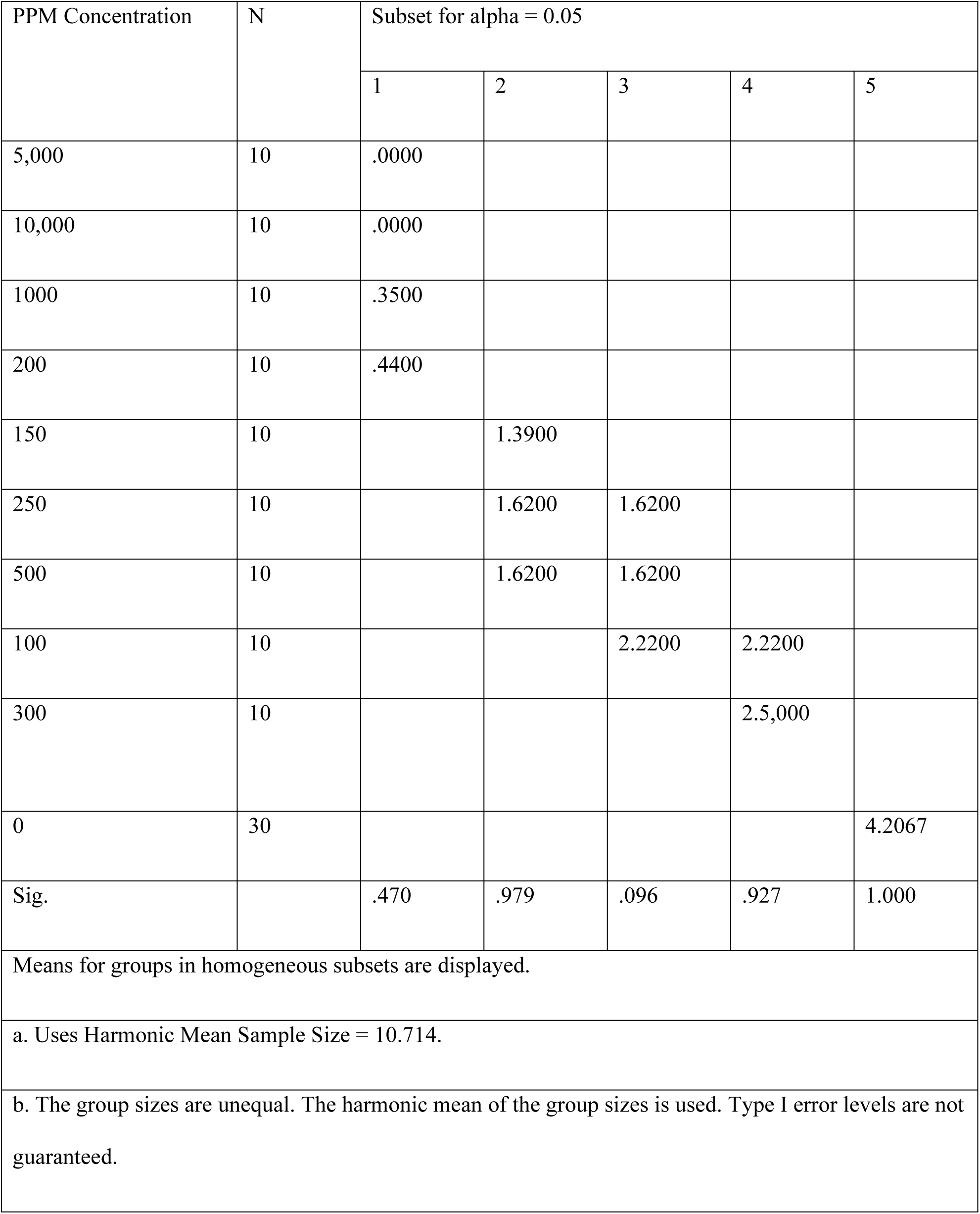
Tukey HSD Analysis of Cymoxanil using SPSS.

## S3 Appendix

**Table 14:** Analysis of Pilarzox using SPSS.

| Average Pilarzox radial growth |  |  |  |  |  |  |  |  |
| --- | --- | --- | --- | --- | --- | --- | --- | --- |
| Radial Growth |  |  |  |  |  |  |  |  |
|  | N | Mean | Std.<br>Deviation | Std.<br>Error | 95% Confidence Interval<br>for Mean |  | Minimum | Maximum |
|  |  |  |  |  | Lower<br>Bound | Upper<br>Bound |  |  |
| 0 | 30 | 4.2067 | .34334 | .06269 | 4.0785 | 4.3349 | 3.30 | 4.70 |
| 100 | 20 | 2.3800 | .54638 | .12217 | 2.1243 | 2.6357 | 1.40 | 3.40 |
| 150 | 20 | 1.4600 | .75561 | .16896 | 1.1064 | 1.8136 | .30 | 2.90 |
| 200 | 20 | .9500 | .55583 | .12429 | .6899 | 1.2101 | .40 | 2.00 |
| 250 | 20 | 1.2250 | .72900 | .16301 | .8838 | 1.5662 | .00 | 2.40 |
| 300 | 20 | 1.5200 | 1.04660 | .23403 | 1.0302 | 2.0098 | .00 | 2.80 |
| 500 | 20 | .8100 | .84161 | .18819 | .4161 | 1.2039 | .00 | 1.90 |
| 1000 | 20 | .1750 | .18317 | .04096 | .0893 | .2607 | .00 | .40 |
| 5,000 | 20 | .0000 | .00000 | .00000 | .0000 | .0000 | .00 | .00 |
| 10,000 | 20 | .0000 | .00000 | .00000 | .0000 | .0000 | .00 | .00 |
| Total | 210 | 1.4124 | 1.46197 | .10089 | 1.2135 | 1.6113 | .00 | 4.70 |

**Table 15:**
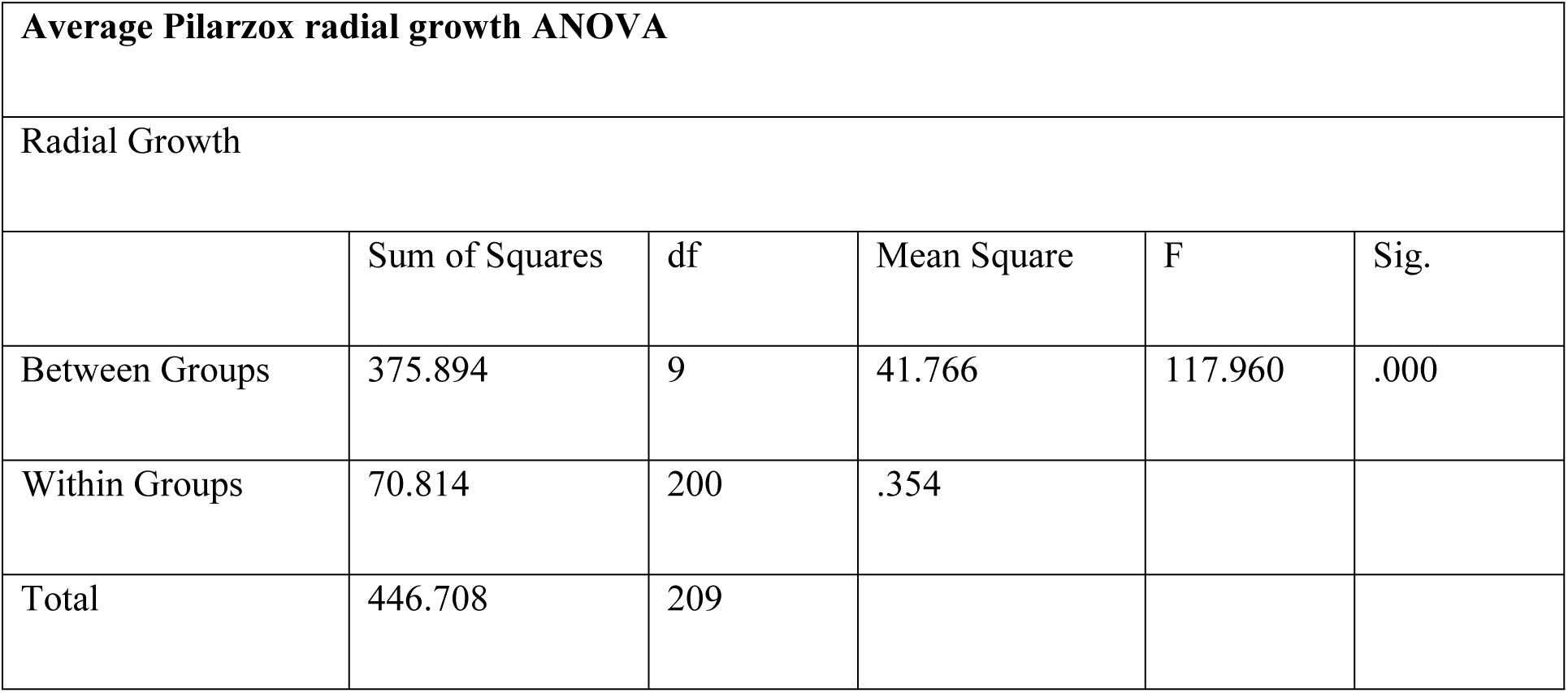
ANOVA Analysis of Pilarzox using SPSS.

| **Average Pilarzox radial growth ANOVA** |  |  |  |  |  |
| --- | --- | --- | --- | --- | --- |
| Radial Growth |  |  |  |  |  |
|  | Sum of Squares | df | Mean Square | F | Sig. |
| Between Groups | 375.894 | 9 | 41.766 | 117.960 | .000 |
| Within Groups | 70.814 | 200 | .354 |  |  |
| Total | 446.708 | 209 |  |  |  |

**Table 16:** Multiple Comparison Analysis of Pilarzox using SPSS.

| Average Pilarzox Radial Growth Multiple Comparisons |  |  |  |  |  |  |
| --- | --- | --- | --- | --- | --- | --- |
| Dependent Variable: Radial Growth |  |  |  |  |  |  |
| Tukey HSD |  |  |  |  |  |  |
| (I)<br>PPM_Concentration | (J)<br>PPM_Concentration | Mean<br>Difference (I-J) | Std.<br>Error | Sig. | 95% Confidence<br>Interval |  |
|  |  |  |  |  | Lower<br>Bound | Upper<br>Bound |
| 0 | 100 | 1.82667* | .17177 | .000 | 1.2770 | 2.3763 |
|  | 150 | 2.74667* | .17177 | .000 | 2.1970 | 3.2963 |
|  | 200 | 3.25667* | .17177 | .000 | 2.7070 | 3.8063 |
|  | 250 | 2.98167* | .17177 | .000 | 2.4320 | 3.5313 |
|  | 300 | 2.68667* | .17177 | .000 | 2.1370 | 3.2363 |
|  | 500 | 3.39667* | .17177 | .000 | 2.8470 | 3.9463 |
|  | 1000 | 4.03167* | .17177 | .000 | 3.4820 | 4.5813 |
|  | 5,000 | 4.20667* | .17177 | .000 | 3.6570 | 4.7563 |
|  | 10,000 | 4.20667* | .17177 | .000 | 3.6570 | 4.7563 |
| 100 | 0 | -1.82667* | .17177 | .000 | -2.3763 | -1.2770 |
|  | 150 | .92000* | .18817 | .000 | .3179 | 1.5221 |
|  | 200 | 1.43000* | .18817 | .000 | .8279 | 2.0321 |
|  | 250 | 1.15500* | .18817 | .000 | .5529 | 1.7571 |
|  | 300 | .86000* | .18817 | .000 | .2579 | 1.4621 |
|  | 500 | 1.57000* | .18817 | .000 | .9679 | 2.1721 |
|  | 1000 | 2.20500* | .18817 | .000 | 1.6029 | 2.8071 |
|  | 5,000 | 2.38000* | .18817 | .000 | 1.7779 | 2.9821 |
|  | 10,000 | 2.38000* | .18817 | .000 | 1.7779 | 2.9821 |
| 150 | 0 | -2.74667* | .17177 | .000 | -3.2963 | -2.1970 |
|  | 100 | -.92000* | .18817 | .000 | -1.5221 | -.3179 |
|  | 200 | .51000 | .18817 | .177 | -.0921 | 1.1121 |
|  | 250 | .23500 | .18817 | .963 | -.3671 | .8371 |
|  | 300 | -.06000 | .18817 | 1.000 | -.6621 | .5421 |
|  | 500 | .65,000* | .18817 | .023 | .0479 | 1.2521 |
|  | 1000 | 1.28500* | .18817 | .000 | .6829 | 1.8871 |
|  | 5,000 | 1.46000* | .18817 | .000 | .8579 | 2.0621 |
|  | 10,000 | 1.46000* | .18817 | .000 | .8579 | 2.0621 |
| 200 | 0 | -3.25667* | .17177 | .000 | -3.8063 | -2.7070 |
|  | 100 | -1.43000* | .18817 | .000 | -2.0321 | -.8279 |
|  | 150 | -.51000 | .18817 | .177 | -1.1121 | .0921 |
|  | 250 | -.27500 | .18817 | .905 | -.8771 | .3271 |
|  | 300 | -.57000 | .18817 | .080 | -1.1721 | .0321 |
|  | 500 | .14000 | .18817 | .999 | -.4621 | .7421 |
|  | 1000 | .77500* | .18817 | .002 | .1729 | 1.3771 |
|  | 5,000 | .95,000* | .18817 | .000 | .3479 | 1.5521 |
|  | 10,000 | .95,000* | .18817 | .000 | .3479 | 1.5521 |
| 250 | 0 | -2.98167* | .17177 | .000 | -3.5313 | -2.4320 |
|  | 100 | -1.15500* | .18817 | .000 | -1.7571 | -.5529 |
|  | 150 | -.23500 | .18817 | .963 | -.8371 | .3671 |
|  | 200 | .27500 | .18817 | .905 | -.3271 | .8771 |
|  | 300 | -.29500 | .18817 | .862 | -.8971 | .3071 |
|  | 500 | .41500 | .18817 | .457 | -.1871 | 1.0171 |
|  | 1000 | 1.05,000* | .18817 | .000 | .4479 | 1.6521 |
|  | 5,000 | 1.22500* | .18817 | .000 | .6229 | 1.8271 |
|  | 10,000 | 1.22500* | .18817 | .000 | .6229 | 1.8271 |
| 300 | 0 | -2.68667* | .17177 | .000 | -3.2363 | -2.1370 |
|  | 100 | -.86000* | .18817 | .000 | -1.4621 | -.2579 |
|  | 150 | .06000 | .18817 | 1.000 | -.5421 | .6621 |
|  | 200 | .57000 | .18817 | .080 | -.0321 | 1.1721 |
|  | 250 | .29500 | .18817 | .862 | -.3071 | .8971 |
|  | 500 | .71000* | .18817 | .008 | .1079 | 1.3121 |
|  | 1000 | 1.34500* | .18817 | .000 | .7429 | 1.9471 |
|  | 5,000 | 1.52000* | .18817 | .000 | .9179 | 2.1221 |
|  | 10,000 | 1.52000* | .18817 | .000 | .9179 | 2.1221 |
| 500 | 0 | -3.39667* | .17177 | .000 | -3.9463 | -2.8470 |
|  | 100 | -1.57000* | .18817 | .000 | -2.1721 | -.9679 |
|  | 150 | -.65,000* | .18817 | .023 | -1.2521 | -.0479 |
|  | 200 | -.14000 | .18817 | .999 | -.7421 | .4621 |
|  | 250 | -.41500 | .18817 | .457 | -1.0171 | .1871 |
|  | 300 | -.71000* | .18817 | .008 | -1.3121 | -.1079 |
|  | 1000 | .63500* | .18817 | .030 | .0329 | 1.2371 |
|  | 5,000 | .81000* | .18817 | .001 | .2079 | 1.4121 |
|  | 10,000 | .81000* | .18817 | .001 | .2079 | 1.4121 |
| 1000 | 0 | -4.03167* | .17177 | .000 | -4.5813 | -3.4820 |
|  | 100 | -2.20500* | .18817 | .000 | -2.8071 | -1.6029 |
|  | 150 | -1.28500* | .18817 | .000 | -1.8871 | -.6829 |
|  | 200 | -.77500* | .18817 | .002 | -1.3771 | -.1729 |
|  | 250 | -1.05,000* | .18817 | .000 | -1.6521 | -.4479 |
|  | 300 | -1.34500* | .18817 | .000 | -1.9471 | -.7429 |
|  | 500 | -.63500* | .18817 | .030 | -1.2371 | -.0329 |
|  | 5,000 | .17500 | .18817 | .995 | -.4271 | .7771 |
|  | 10,000 | .17500 | .18817 | .995 | -.4271 | .7771 |
| 5,000 | 0 | -4.20667* | .17177 | .000 | -4.7563 | -3.6570 |
|  | 100 | -2.38000* | .18817 | .000 | -2.9821 | -1.7779 |
|  | 150 | -1.46000* | .18817 | .000 | -2.0621 | -.8579 |
|  | 200 | -.95,000* | .18817 | .000 | -1.5521 | -.3479 |
|  | 250 | -1.22500* | .18817 | .000 | -1.8271 | -.6229 |
|  | 300 | -1.52000* | .18817 | .000 | -2.1221 | -.9179 |
|  | 500 | -.81000* | .18817 | .001 | -1.4121 | -.2079 |
|  | 1000 | -.17500 | .18817 | .995 | -.7771 | .4271 |
|  | 10,000 | .00000 | .18817 | 1.000 | -.6021 | .6021 |
| 10,000 | 0 | -4.20667* | .17177 | .000 | -4.7563 | -3.6570 |
|  | 100 | -2.38000* | .18817 | .000 | -2.9821 | -1.7779 |
|  | 150 | -1.46000* | .18817 | .000 | -2.0621 | -.8579 |
|  | 200 | -.95,000* | .18817 | .000 | -1.5521 | -.3479 |
|  | 250 | -1.22500* | .18817 | .000 | -1.8271 | -.6229 |
|  | 300 | -1.52000* | .18817 | .000 | -2.1221 | -.9179 |
|  | 500 | -.81000* | .18817 | .001 | -1.4121 | -.2079 |
|  | 1000 | -.17500 | .18817 | .995 | -.7771 | .4271 |
|  | 5,000 | .00000 | .18817 | 1.000 | -.6021 | .6021 |

\*. The mean difference is significant at the 0.05 level.

**Table 17:**
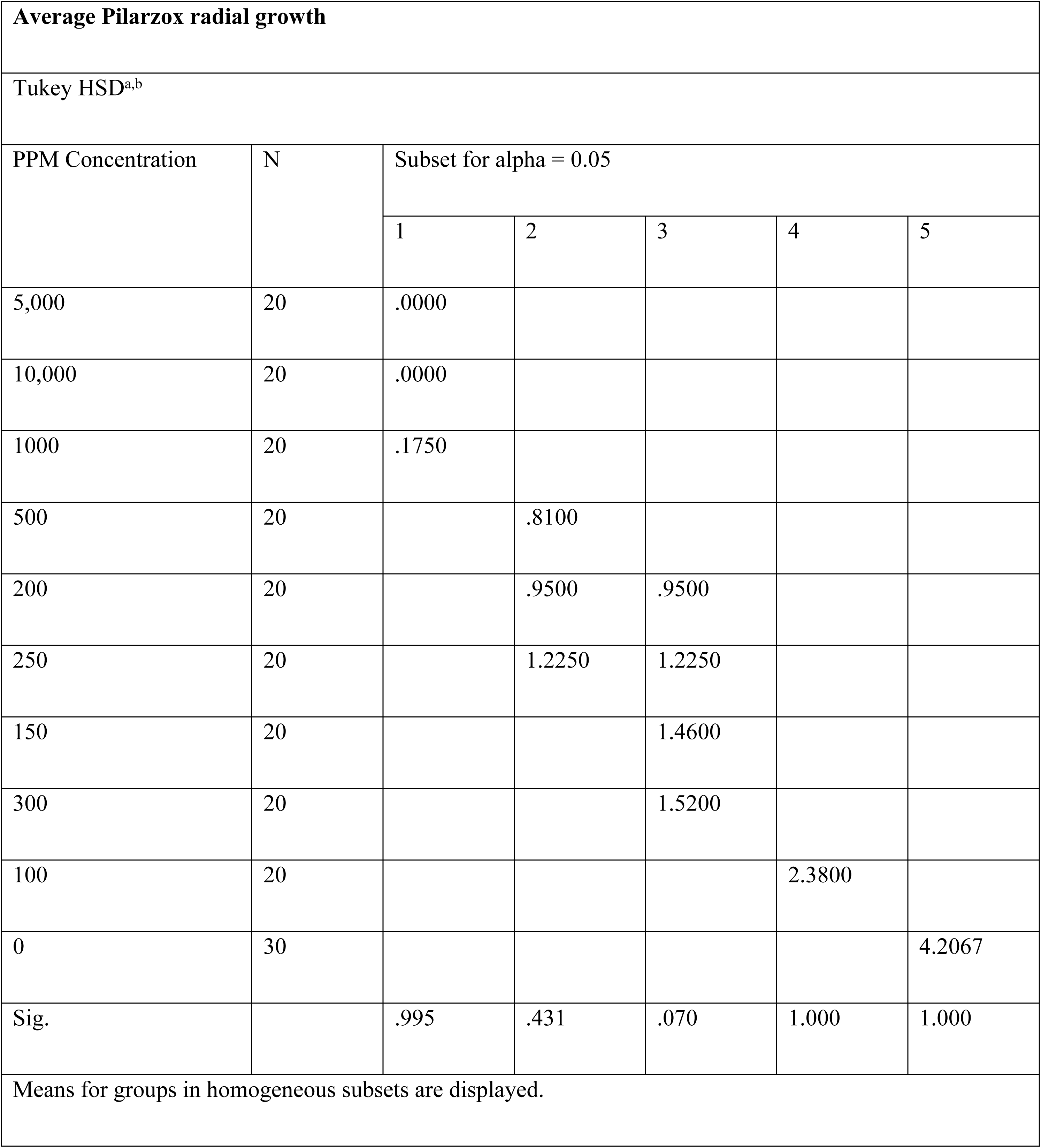
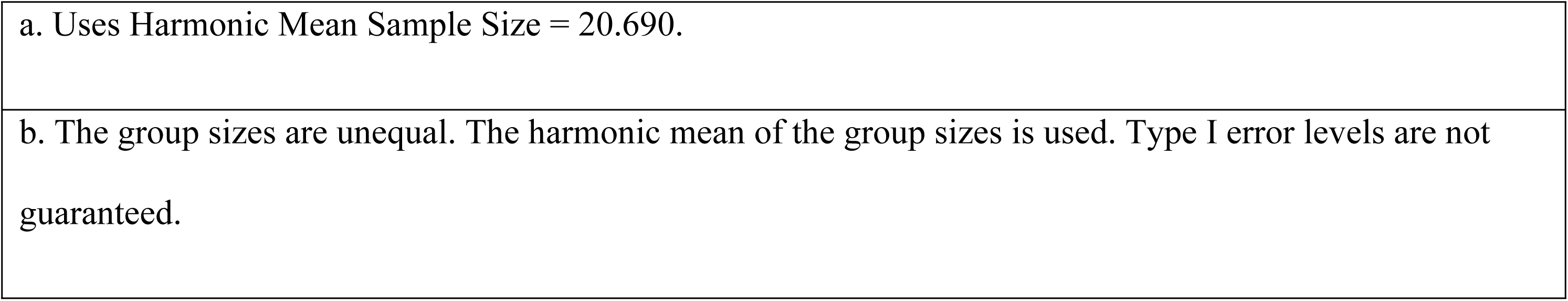
Tukey HSD Analysis of Pilarzox using SPSS.

| Average Pilarzox radial growth |  |  |  |  |  |  |
| --- | --- | --- | --- | --- | --- | --- |
| Tukey HSD<sup>a,b</sup> |  |  |  |  |  |  |
| PPM Concentration | N | Subset for alpha = 0.05 |  |  |  |  |
|  |  | 1 | 2 | 3 | 4 | 5 |
| 5,000 | 20 | .0000 |  |  |  |  |
| 10,000 | 20 | .0000 |  |  |  |  |
| 1000 | 20 | .1750 |  |  |  |  |
| 500 | 20 |  | .8100 |  |  |  |
| 200 | 20 |  | .9500 | .9500 |  |  |
| 250 | 20 |  | 1.2250 | 1.2250 |  |  |
| 150 | 20 |  |  | 1.4600 |  |  |
| 300 | 20 |  |  | 1.5200 |  |  |
| 100 | 20 |  |  |  | 2.3800 |  |
| 0 | 30 |  |  |  |  | 4.2067 |
| Sig. |  | .995 | .431 | .070 | 1.000 | 1.000 |
| Means for groups in homogeneous subsets are displayed. |  |  |  |  |  |  |

|  |
| --- |
| a. Uses Harmonic Mean Sample Size = 20.690. |
| b. The group sizes are unequal. The harmonic mean of the group sizes is used. Type I error levels are not guaranteed. |

## S4 Appendix Turmeric results

**Table 19:** Analysis of Turmeric using SPSS.

| **Turmeric** |  |  |  |  |  |  |  |  |
| --- | --- | --- | --- | --- | --- | --- | --- | --- |
| Radial Growth |  |  |  |  |  |  |  |  |
|  | N | Mean | Std.<br>Deviation | Std.<br>Error | 95% Confidence<br>Interval for Mean |  | Minimum | Maximum |
|  |  |  |  |  | Lower<br>Bound | Upper<br>Bound |  |  |
| 0 | 40 | 4.1100 | .37059 | .05859 | 3.9915 | 4.2285 | 3.20 | 4.70 |
| 100 | 10 | 3.6500 | .42753 | .13520 | 3.3442 | 3.9558 | 2.90 | 4.20 |
| 150 | 10 | 3.8400 | .33400 | .10562 | 3.6011 | 4.0789 | 3.50 | 4.50 |
| 200 | 10 | 3.4900 | .80753 | .25536 | 2.9123 | 4.0677 | 2.00 | 4.20 |
| 250 | 10 | 3.7000 | .32998 | .10435 | 3.4639 | 3.9361 | 3.20 | 4.20 |
| 300 | 10 | 3.8100 | .15239 | .04819 | 3.7010 | 3.9190 | 3.50 | 4.00 |
| 500 | 10 | 3.7900 | .27669 | .08750 | 3.5921 | 3.9879 | 3.20 | 4.20 |
| 1000 | 10 | 3.7800 | .19322 | .06110 | 3.6418 | 3.9182 | 3.50 | 4.20 |
| 5,000 | 10 | 3.6000 | .20548 | .06498 | 3.4530 | 3.7470 | 3.20 | 3.90 |
| 10,000 | 10 | 3.3500 | .27183 | .08596 | 3.1555 | 3.5445 | 3.00 | 3.80 |
| Total | 130 | 3.8038 | .43765 | .03838 | 3.7279 | 3.8798 | 2.00 | 4.70 |

**Table 20:** ANOVA Analysis of Turmeric using SPSS.

| **ANOVA** |  |  |  |  |  |
| --- | --- | --- | --- | --- | --- |
| Radial Growth |  |  |  |  |  |
|  | Sum of Squares | df | Mean Square | F | Sig. |
| Between Groups | 7.575 | 9 | .842 | 5.895 | .000 |
| Within Groups | 17.133 | 120 | .143 |  |  |
| Total | 24.708 | 129 |  |  |  |

**Table 21:**
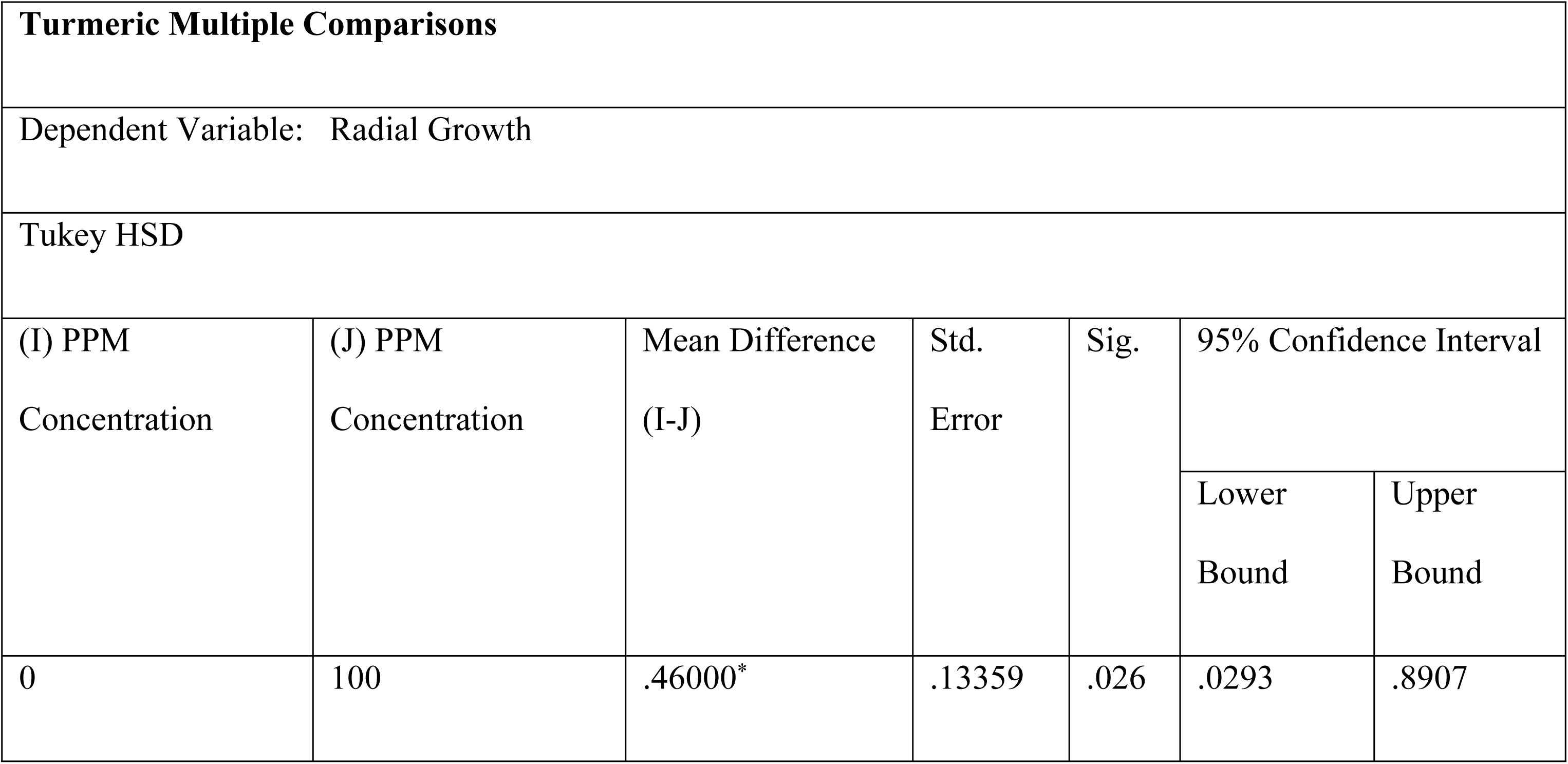
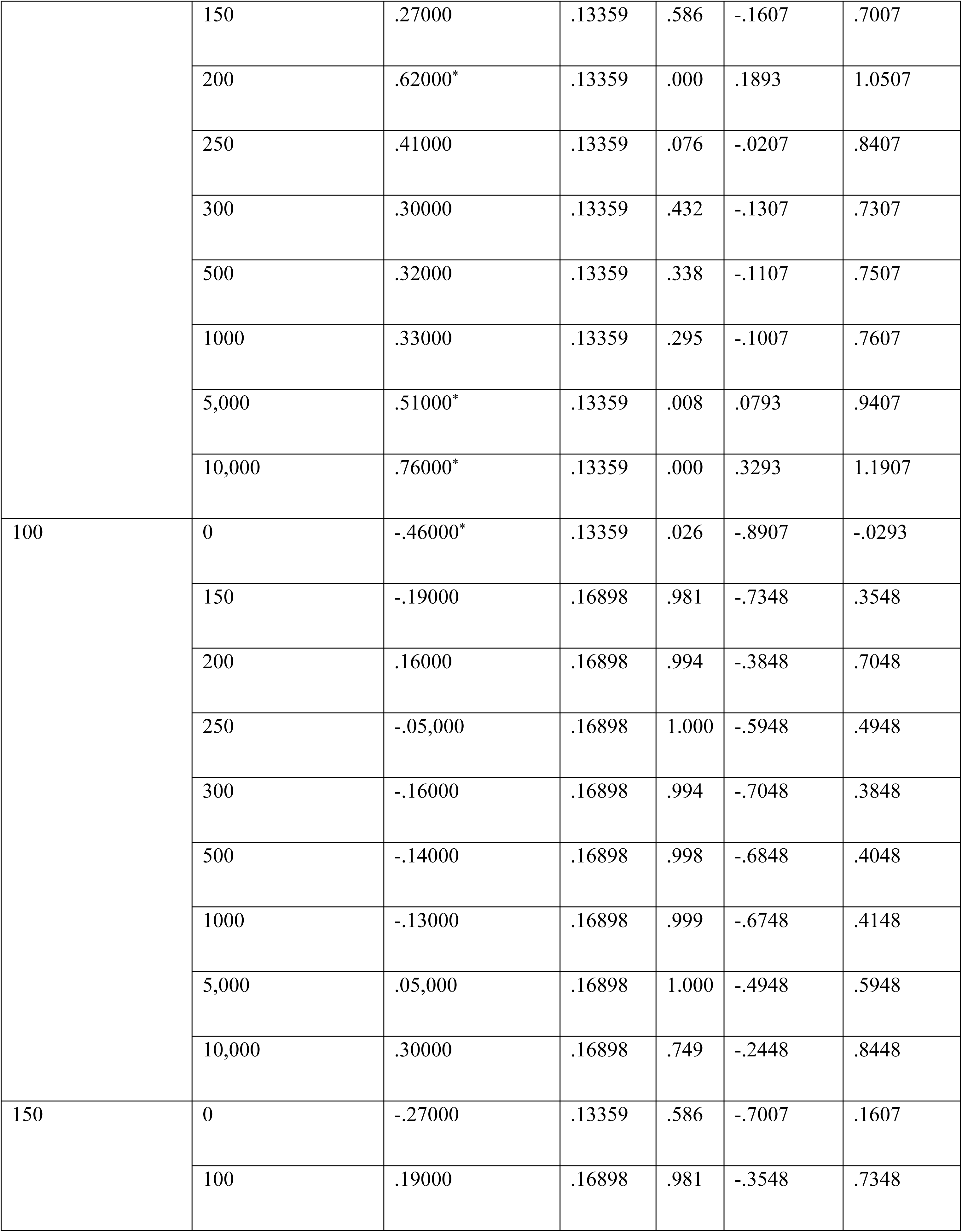
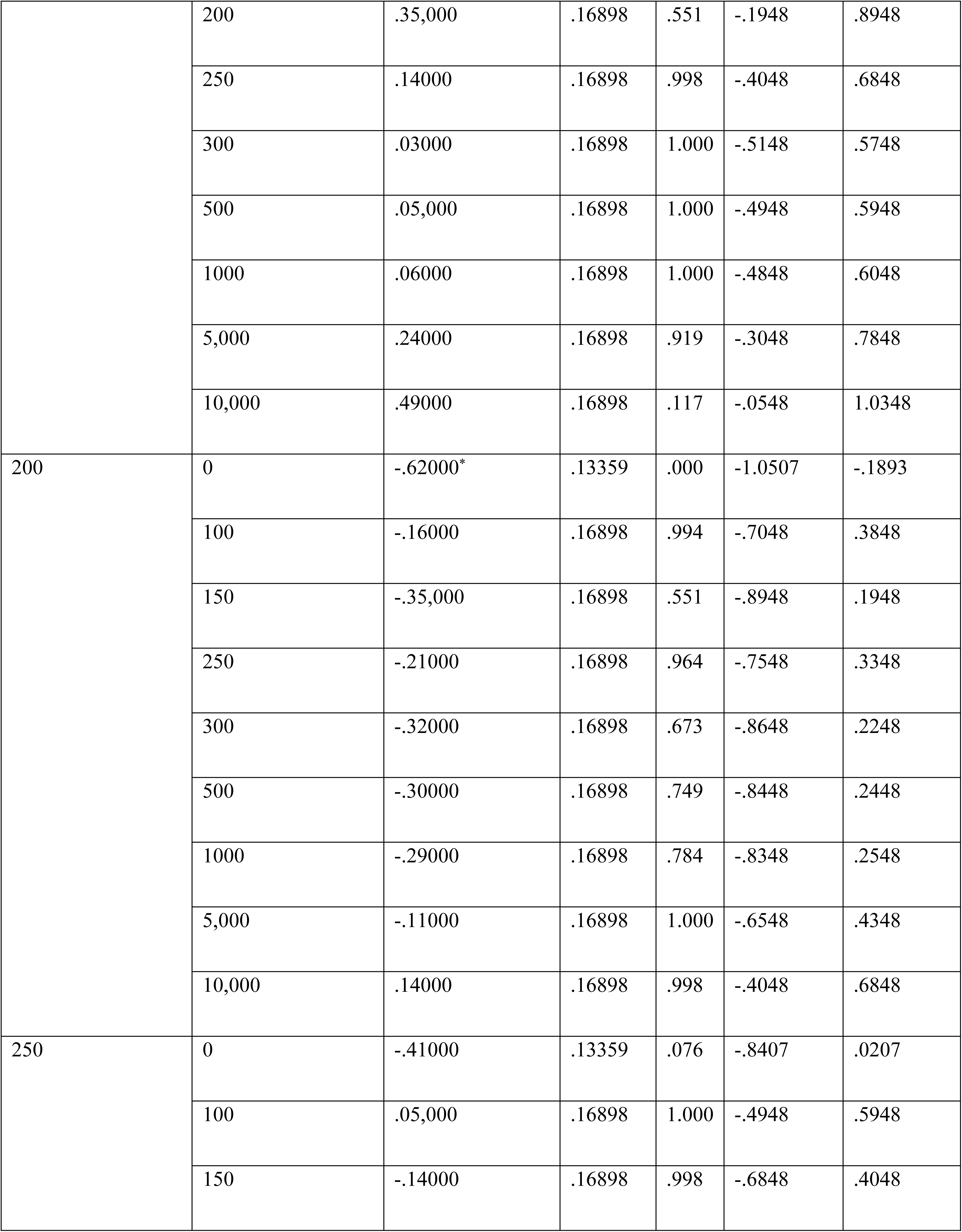
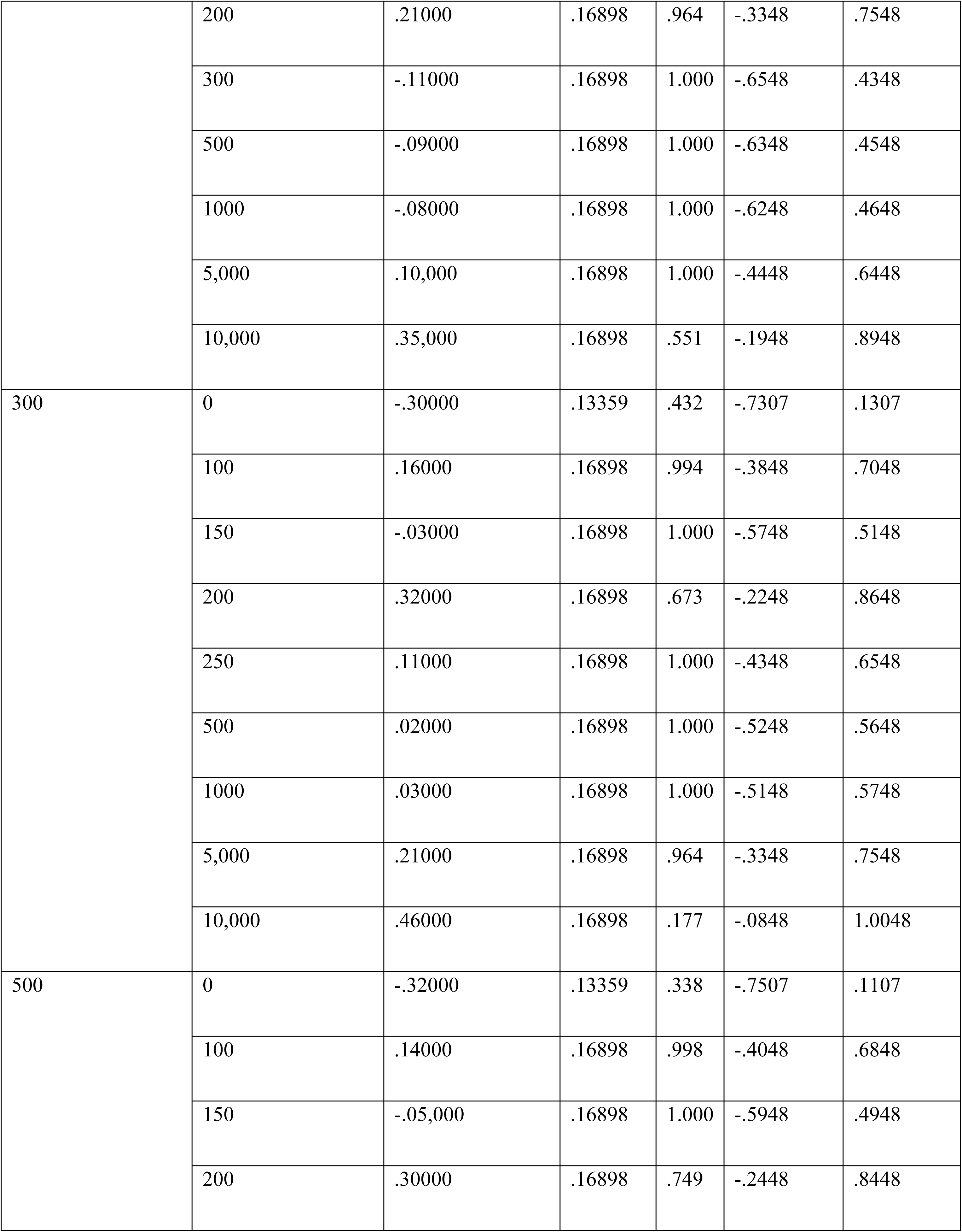
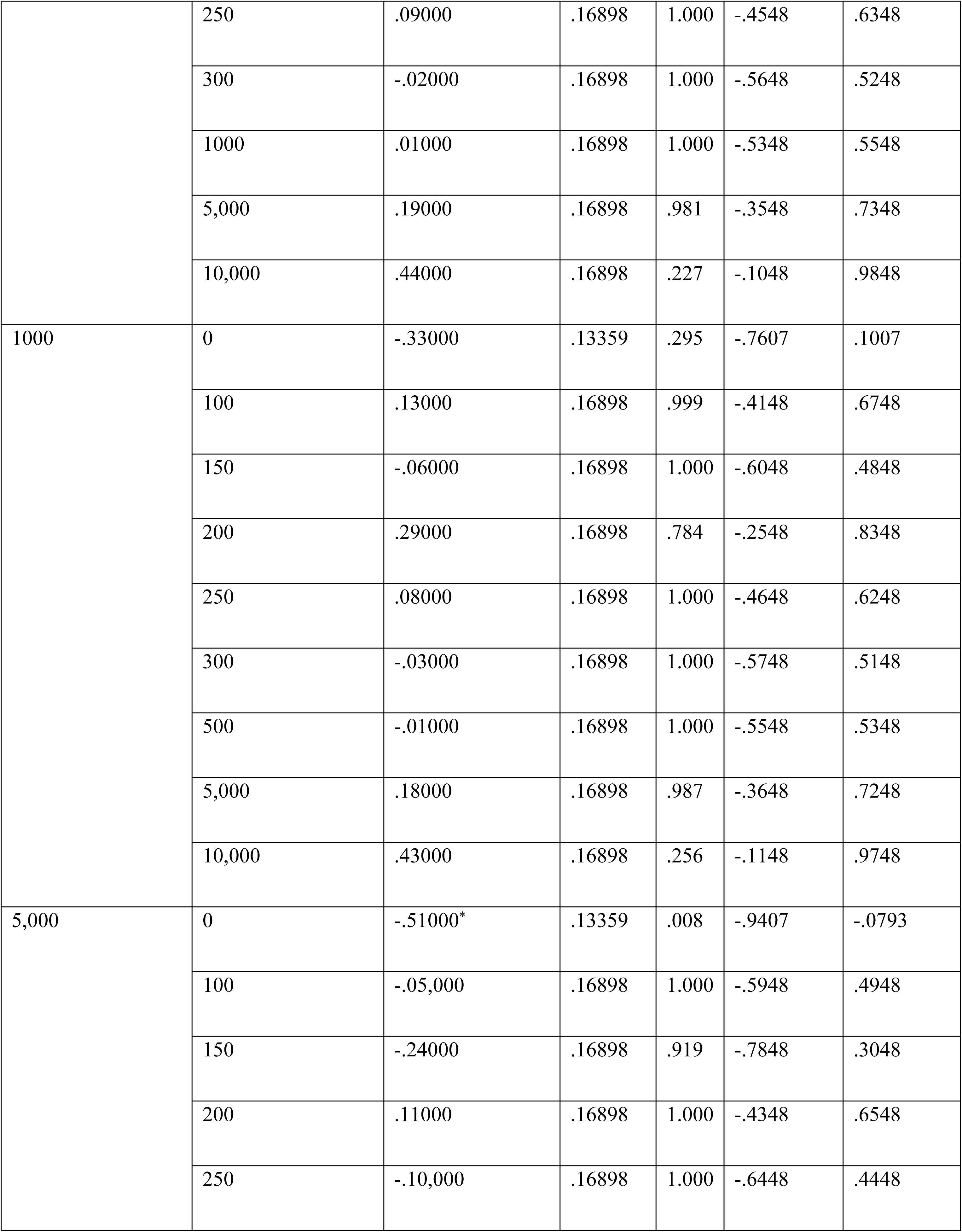
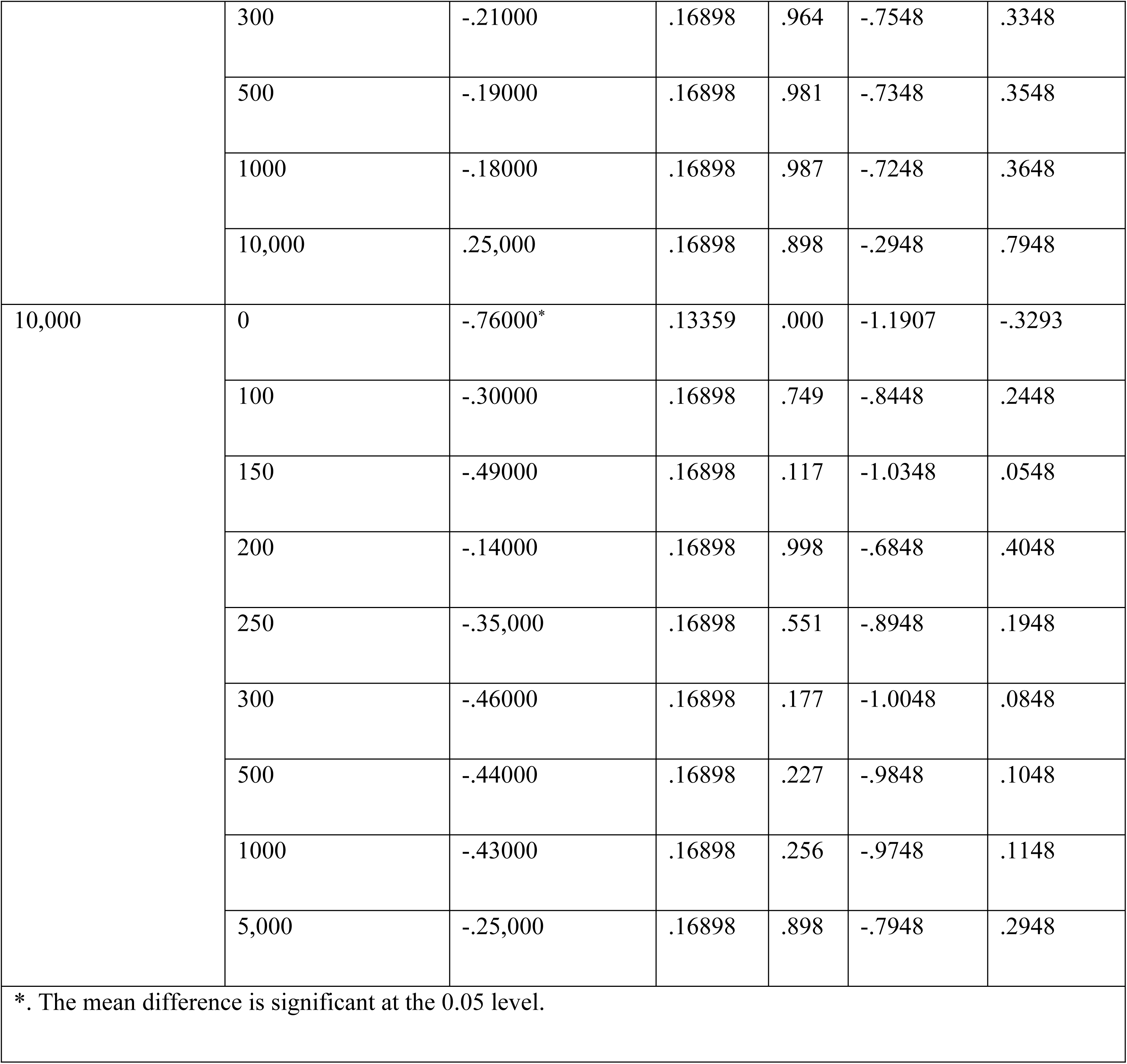
Multiple Comparison Analysis of Turmeric using SPSS.

**Table 22:** Tukey HSD Analysis of Turmeric using SPSS.

| **Turmeric Radial Growth** |  |  |  |
| --- | --- | --- | --- |
| Tukey HSD<sup>a,b</sup> |  |  |  |
| PPM_Concentration | N | Subset for alpha = 0.05 |  |
|  |  | 1 | 2 |
| 10,000 | 10 | 3.3500 |  |
| 200 | 10 | 3.4900 |  |
| 5,000 | 10 | 3.6000 | 3.6000 |
| 100 | 10 | 3.6500 | 3.6500 |
| 250 | 10 | 3.7000 | 3.7000 |
| 1000 | 10 | 3.7800 | 3.7800 |
| 500 | 10 | 3.7900 | 3.7900 |
| 300 | 10 | 3.8100 | 3.8100 |
| 150 | 10 | 3.8400 | 3.8400 |
| 0 | 40 |  | 4.1100 |
| Sig. |  | .088 | .063 |
| Means for groups in homogeneous subsets are displayed. |  |  |  |
| a. Uses Harmonic Mean Sample Size = 10.811. |  |  |  |
| b. The group sizes are unequal. The harmonic mean of the group sizes is used. Type I error levels are not guaranteed. |  |  |  |

**Figure 1.**
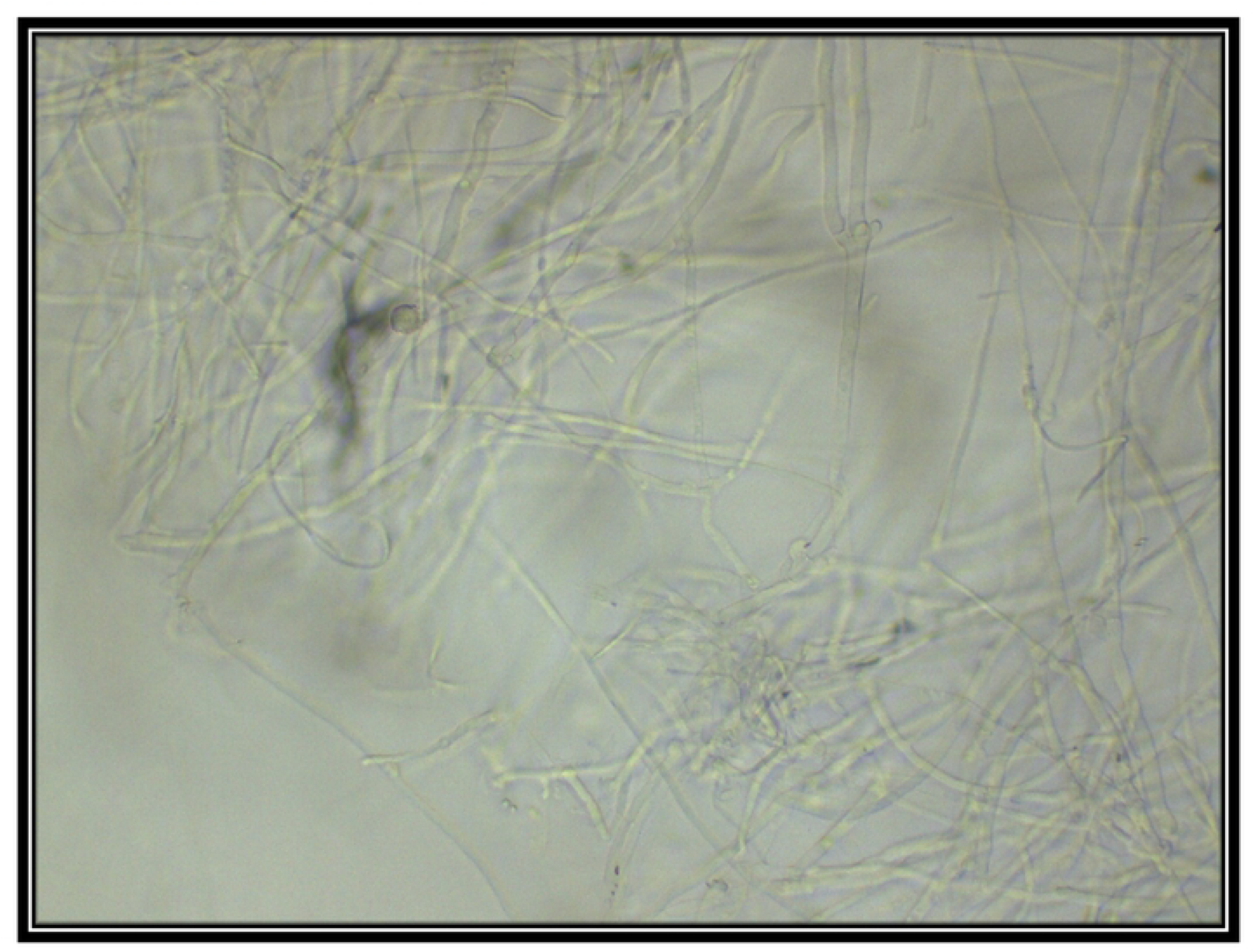
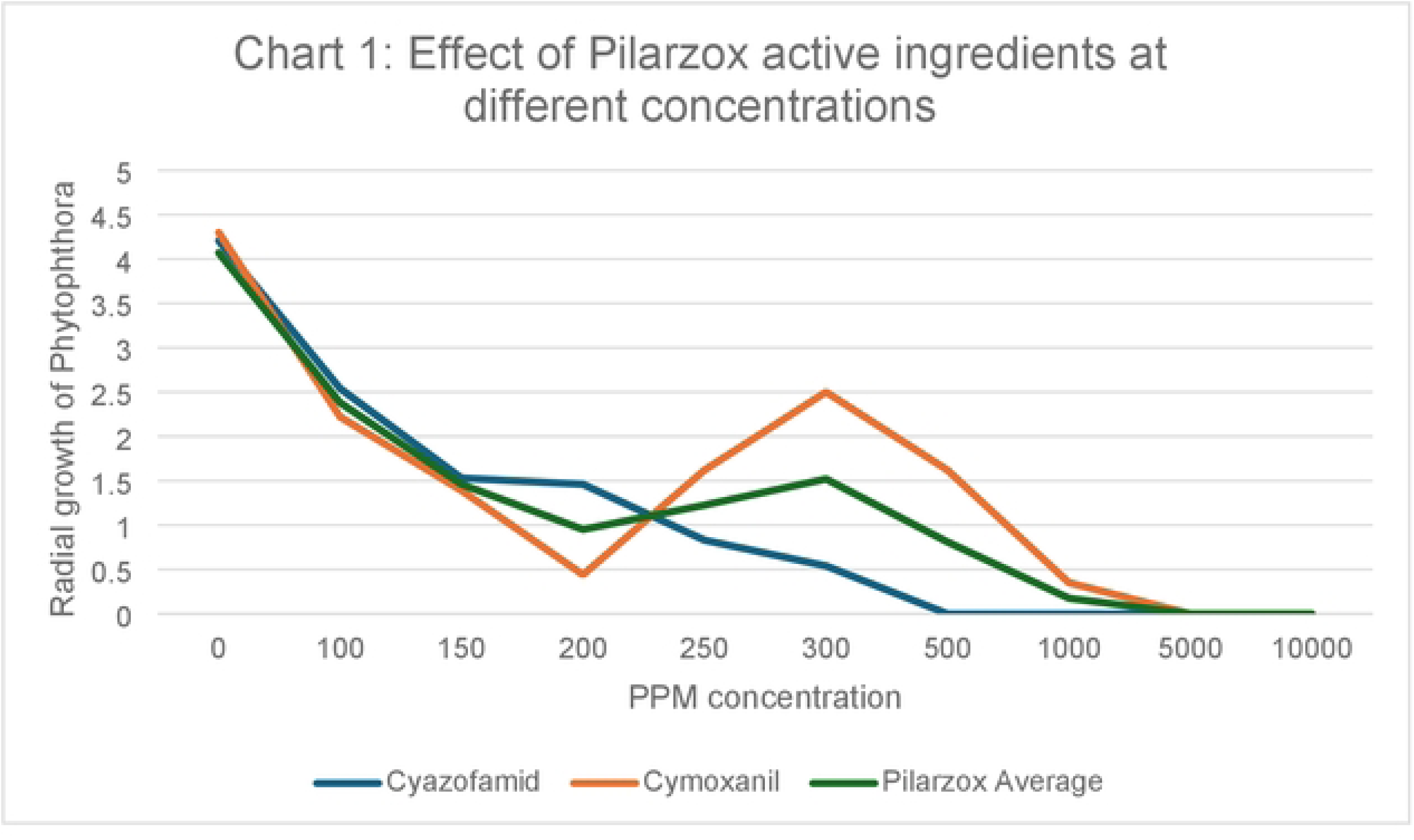
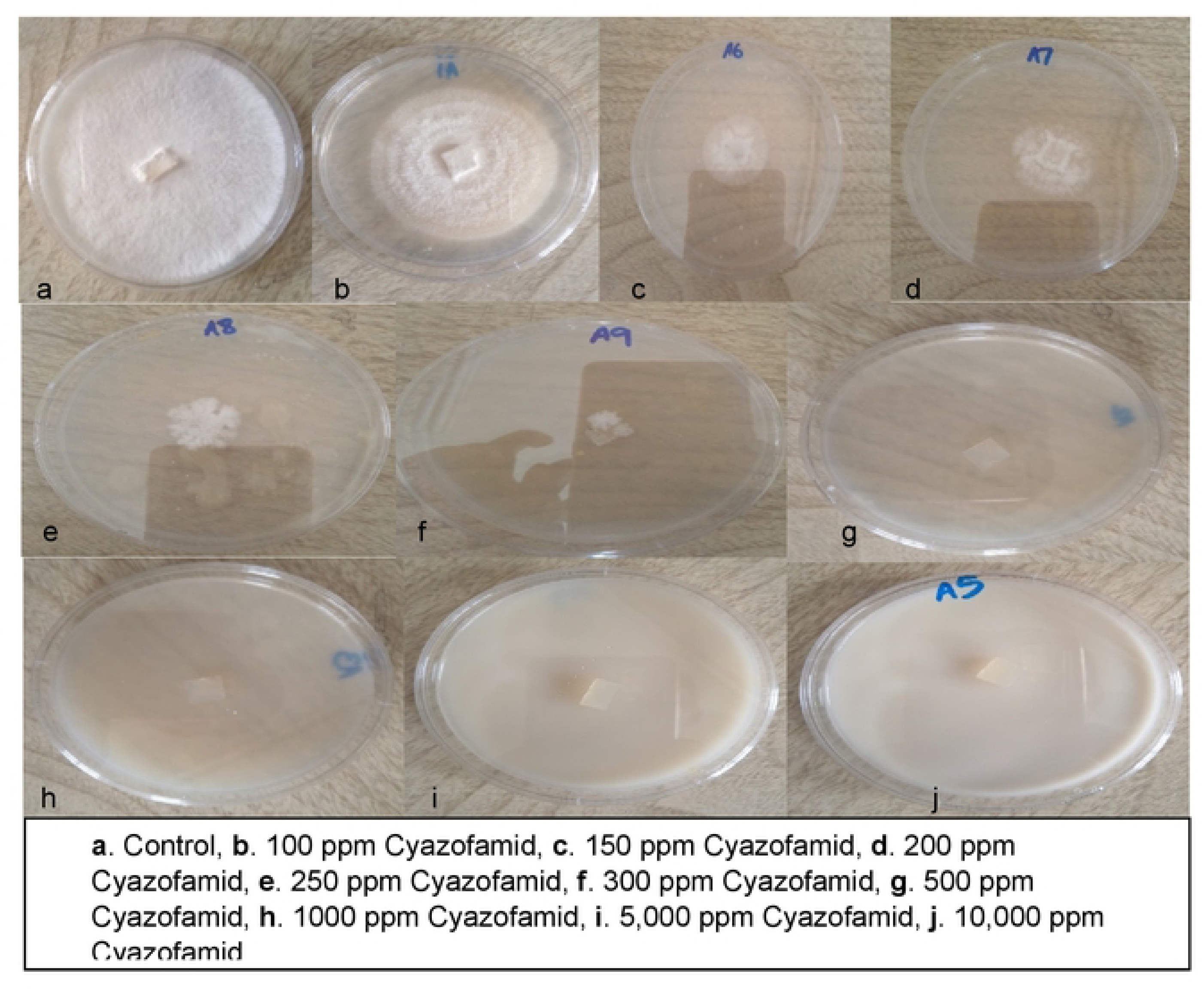
Microscopic view of Phytophlhora sp. Isolate.

**Figure 2.**
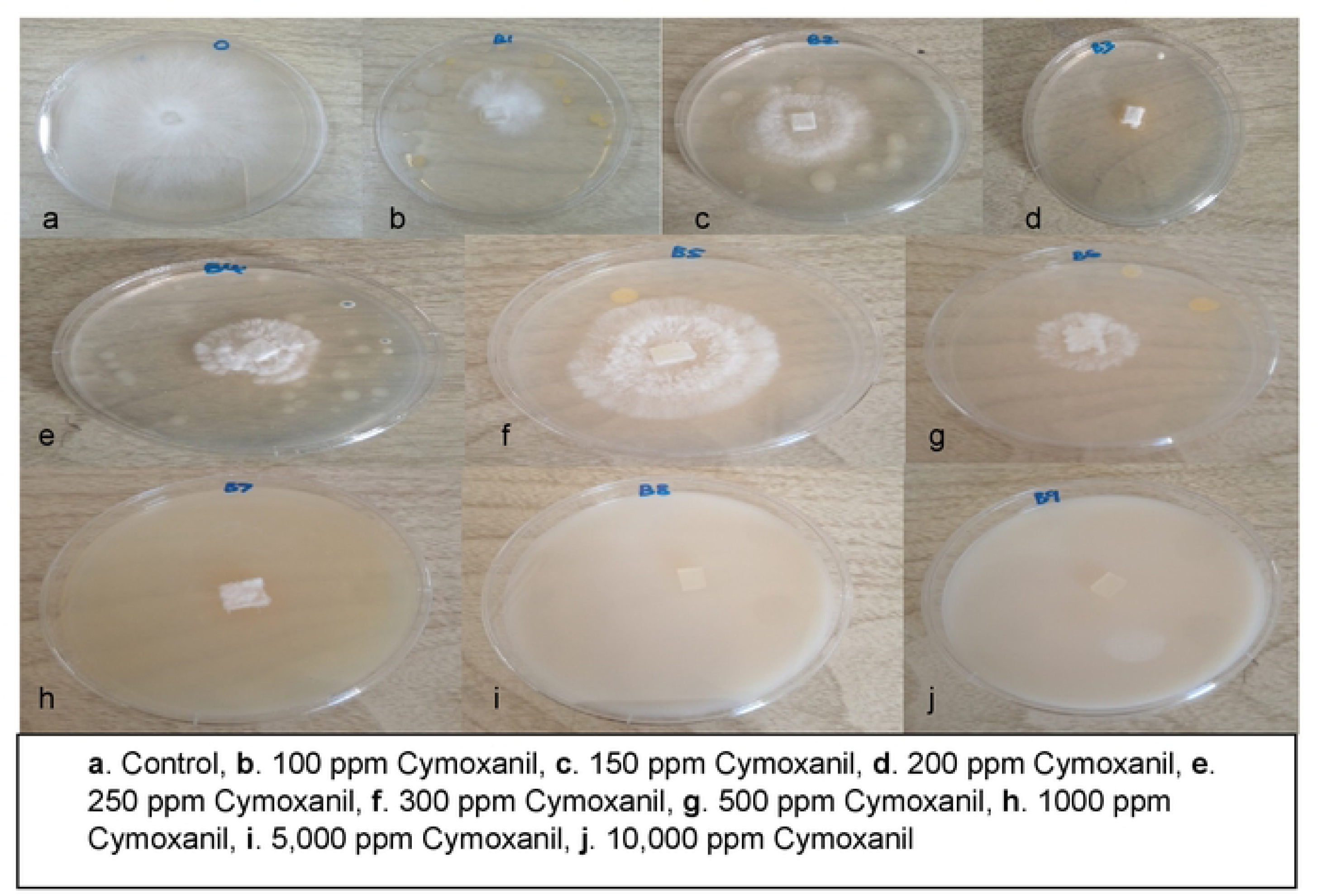
Growth of Phytophthora on PDA with active ingredient Cyazofamid at different concentrations.

**Figure 3.**
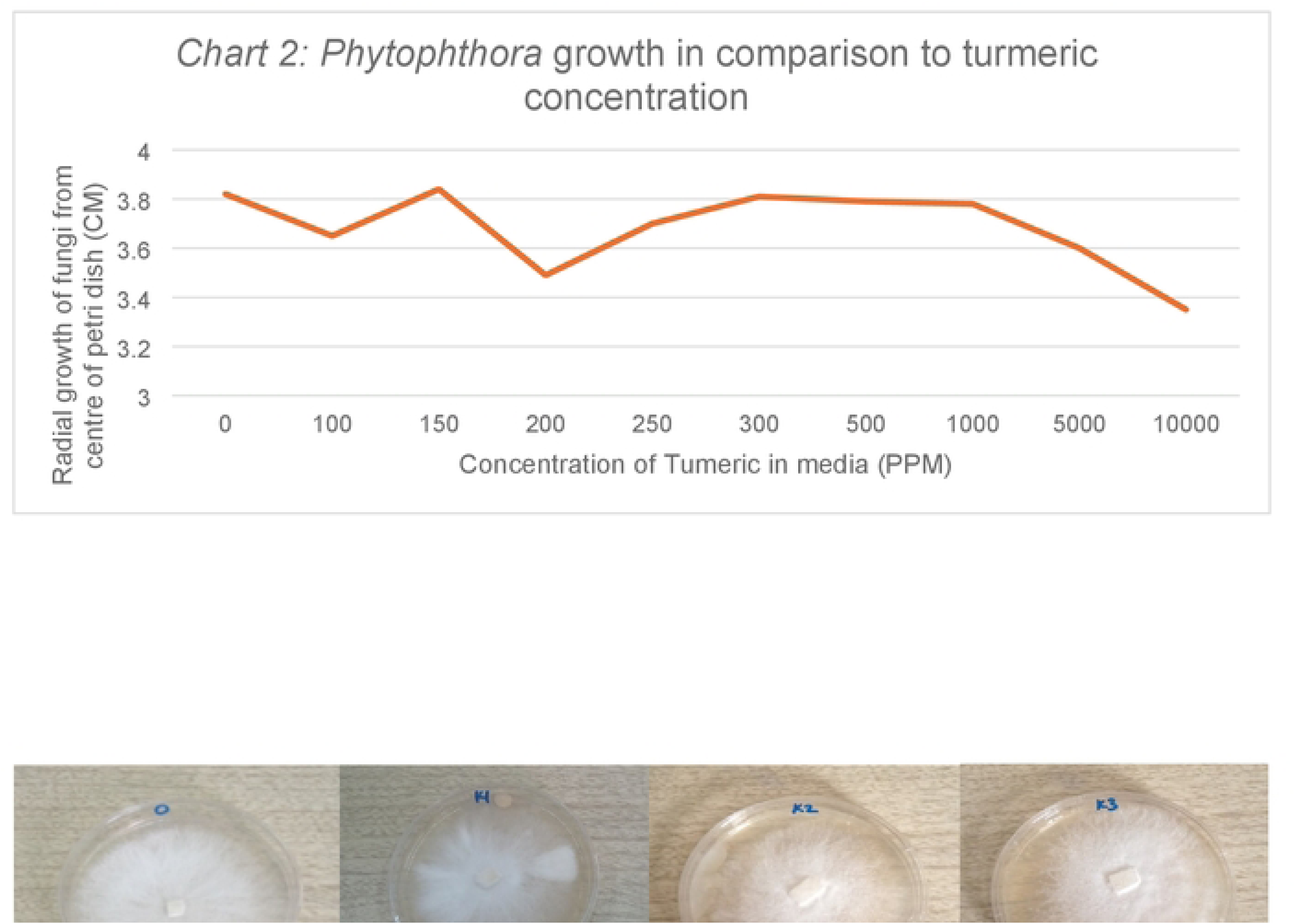
Growth of Phytophthora on PDA with active ingredient Cymoxanil at different concentrations.

